# Benthic diatoms navigate shear flows via hydrodynamic rolling and active gliding

**DOI:** 10.64898/2026.05.09.721219

**Authors:** Bohan Wang, Sheng Ding, Weiquan Jiang, Xinlei Guo, Rui Han, Li Zeng, Jianhua Wang, T.J. Pedley

## Abstract

Navigating fluid flow is a fundamental challenge for microbial life across diverse aquatic environments. While rheotaxis in swimming microorganisms has been extensively studied, it remains unresolved whether near-bed shear merely perturbs gliding motility or instead provides directional cues for active navigation on surfaces. Here we show that the benthic diatom *Navicula cryptocephala* utilises a purely mechanical strategy to achieve downstream rheotaxis and anisotropic spreading on submerged surfaces. Single-cell ellipsoidal tracking reveals a direction-dependent angular response that reorients gliding cells towards the downstream direction. Using interference reflection microscopy, we further reveal that shear induces rolling of obliquely gliding cells, laterally shifting the cell–substrate contact site. This shift renders raphe-based propulsion non-collinear with substrate friction, generating a downstream-restoring yaw torque. Crucially, our results rule out alternative explanations based on longitudinal shifts of the raphe contact site or direct hydrodynamic yaw torque. A minimal stochastic model confirms that this mechanical reorientation alone is sufficient to reproduce the observed drift and diffusion patterns, without invoking either orientation-dependent switching between motility states or orientation-dependent dwell times of those states. Our findings uncover a mechanism by which ambient shear is converted into directional guidance for active surface motility, providing new insights into microbial transport, retention, and resilience on submerged surfaces.

## 1 Introduction

Surface-associated motility shapes how microorganisms colonise, exploit, and persist on surfaces [1–12]. Among the most abundant surface gliders in sunlit submerged habitats, pennate raphid diatoms contribute substantially to a wide range of ecological processes, including primary production, nutrient cycling, and biofilm formation [13–23]. These cells are generally thought to move by raphe-based gliding, driven by mucilage secretion and intracellular actin–myosin activity without extracellular appendages or large body deformations [24–27]. In quiescent media, their motility depends strongly on cell geometry, raphe morphology and substrate interactions [28–32].

Yet benthic diatoms rarely experience completely still water in nature. Instead, they inhabit surfaces strongly exposed to currents, tides and waves, where near-bed shear flow acts continuously on surface-attached cells [33– 37]. Whether such shear flow merely perturbs surface gliding or instead provides directional information that cells can exploit remains unresolved [38]. This question is non-trivial because most motile diatoms are elongated cells that glide predominantly along their longitudinal axis [31, 39], so near-bed shear flow should impose strongly orientation-dependent forcing. At the same time, unlike suspended phytoplankton, gliding diatoms are mechanically coupled to the substrate through adhesion, traction, friction, and other forces. Their response therefore cannot be inferred from either the locomotion of free-swimming microorganisms [40–50] or from gliding diatoms in quiescent media [30–32, 51].

Here we show that the benthic diatom *Navicula cryptocephala* actively navigates shear flow through downstream rheotaxis accompanied by strongly anisotropic diffusion. Combining single-cell tracking, interference reflection microscopy (IRM), hydrodynamic simulations and stochastic modelling, we find that near-bed shear flow rolls obliquely gliding cells and laterally shifts the cell–substrate contact site. This shift makes raphe-based propulsion non-collinear with substrate resistance, generating a restoring yaw torque that aligns cells downstream despite the opposing hydrodynamic yaw torque. These results identify a physical mechanism by which ambient shear flow can guide active migration on surfaces, with implications for microbial transport, retention and resilience on submerged surfaces.

## 2 Results

### 2.1 Most diatoms remain attached and motile under shear flow

The freshwater pennate raphid diatom *Navicula cryptocephala* is an elongated, bilaterally symmetric cell, with two nearly straight raphes running along one valve surface and separated by a central nodule (Fig. 1a). Like other raphid diatoms, it adheres to the substrate by secreting extracellular polymeric substances (EPS) through the raphe system and glides using a raphe-based motility machinery (Fig. 1a).

**Fig. 1.**
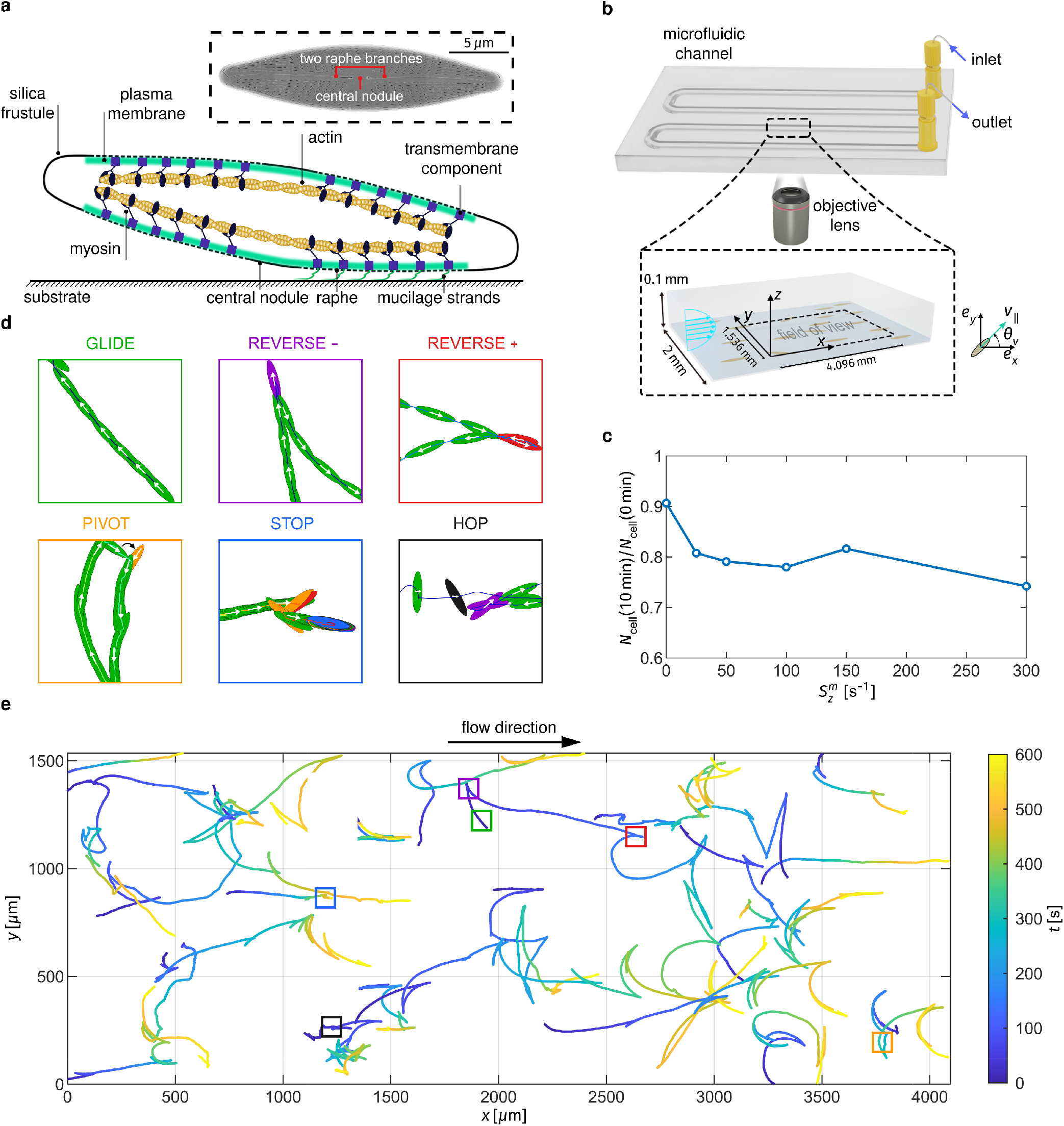
Single-cell tracking of surface-associated gliding under controlled near-bed shear flow. **a**, Schematic side view of a *N. cryptocephala* cell attached to a flat surface and gliding via the raphe-associated actin–myosin motility system. *Inset*: scanning electron microscopy of a single silica frustule, showing the elongated bilaterally symmetric cell shape and the two nearly straight raphes separated by a central nodule. **b**, Microfluidic assay for quantifying gliding motility under imposed shear. Cells were allowed to settle on the bottom wall of a microchannel (width, 2 mm; height, 0.1 mm), after which a cell-free flow of controlled flow rate was applied to generate an approximately uniform vertical bed shear rate over the central analysis region. Trajectories were recorded from the channel bottom using an inverted microscope with a 4× objective. **c**, Fraction of motile cells remaining in the field of view after 10 min of flushing, normalised by the initial number of motile cells, as a function of imposed bed shear rate. Most cells remained attached and motile over the tested range of bed shear rates. **d**, Representative trajectory segments illustrating the five motility states identified from instantaneous gliding speed and angular velocity: GLIDE, REVERSE, STOP, PIVOT and HOP. For REVERSE, the two annotations REVERSE+ and REVERSE-distinguish whether the initially defined positive end becomes the leading or trailing end after reversal, but both belong to the same motility state. Cell outlines are shown at 3 s intervals. **e**, Representative cell trajectories in a field of view of 4.096 *×* 1.536 mm^2^ at 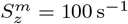, coloured by time. Boxes mark the trajectory segments shown in **d**.

To examine how near-bed shear flow influences surface-associated motility, we allowed a sparse population of cells to settle on the glass bottom of a microfluidic channel before imposing a cell-free medium flow at constant flow rate (Fig. 1b; Methods). This generated an approximately uniform vertical shear rate at the bed, while the lateral shear rate remained negligible over the central 1.536 mm-wide region of the bottom wall, away from the sidewalls, that was used for trajectory analysis (Extended Data Fig. 1; Methods). We tested five non-zero bed shear rates, 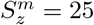, 50, 100, 150 and 300 s^−1^, together with a no-shear control. These values span hydrodynamic conditions commonly encountered in rivers, estuaries, intertidal flats, and open channels [52–55]. Across this range, most cells remained attached and motile, with the total number of motile diatoms in the field of view decreasing by less than 30% after 10 min of flushing (Fig. 1c; Supplementary Video 1). Thus, within an ecologically relevant hydrodynamic range, near-bed shear flow does not simply scour away motile cells, but leaves a substantial surface-associated population capable of gliding on the substrate [36, 37].

*N. cryptocephala* glides in a jerky manner, with fluctuations in both speed and orientation, and a mean gliding speed of approximately 3.8 *µ*m s^−1^ in still water (Fig. 1d, e; Extended Data Fig. 2; Supplementary Video 2). Based on instantaneous gliding speed and angular velocity, we classified tracked segments into five motility states. Four of these—GLIDE, REVERSE, STOP and PIVOT—correspond to the stereotyped states previously identified in still water [31]. GLIDE, the dominant state, is characterised by persistent motion with little directional change and accounts for at least 70% of all trajectory points across all bed shear rates (Extended Data Fig. 3). REVERSE denotes an exchange of the leading and trailing ends and is the second most frequent state, with important consequences for phototaxis and chemotaxis [3, 56, 57]. For classification, we defined the leading end during the first GLIDE event as the positive end and annotated each REVERSE according to whether this end became the leading (+) or trailing (−) end. STOP denotes transient arrest and accounts for less than 10% of trajectory points across all bed shear rates, whereas PIVOT denotes a sharp reorientation [31] and accounts for less than 4%. With an imposed flow, we additionally observed a fifth state, HOP, in which a cell briefly left the substrate but reattached downstream within the field of view; this state was rare, accounting for less than 0.1% of trajectory points. This state-based description provides a framework for resolving how near-bed shear flow alters gliding behaviour and for identifying the dominant factor responsible for effective transport.

### 2.2 Shear-driven cell reorientation underlies downstream rheotaxis and anisotropic diffusion

High-precision bright-field tracking revealed clear downstream rheotaxis in surface-attached cells under shear flow (see Supplementary Videos 2–5 for the extracted trajectories at different bed shear rates): the mean streamwise drift, 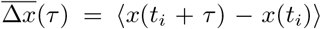, increased monotonically with bed shear rate 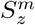. By contrast, the mean transverse drift, 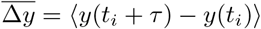, remained close to zero across the full range of 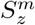 (Fig. 2a). Excluding HOP segments yielded nearly identical drift statistics (Extended Data Fig. 4), indicating that the downstream transport is not a consequence of intermittent scour and passive advection.

**Fig. 2.**
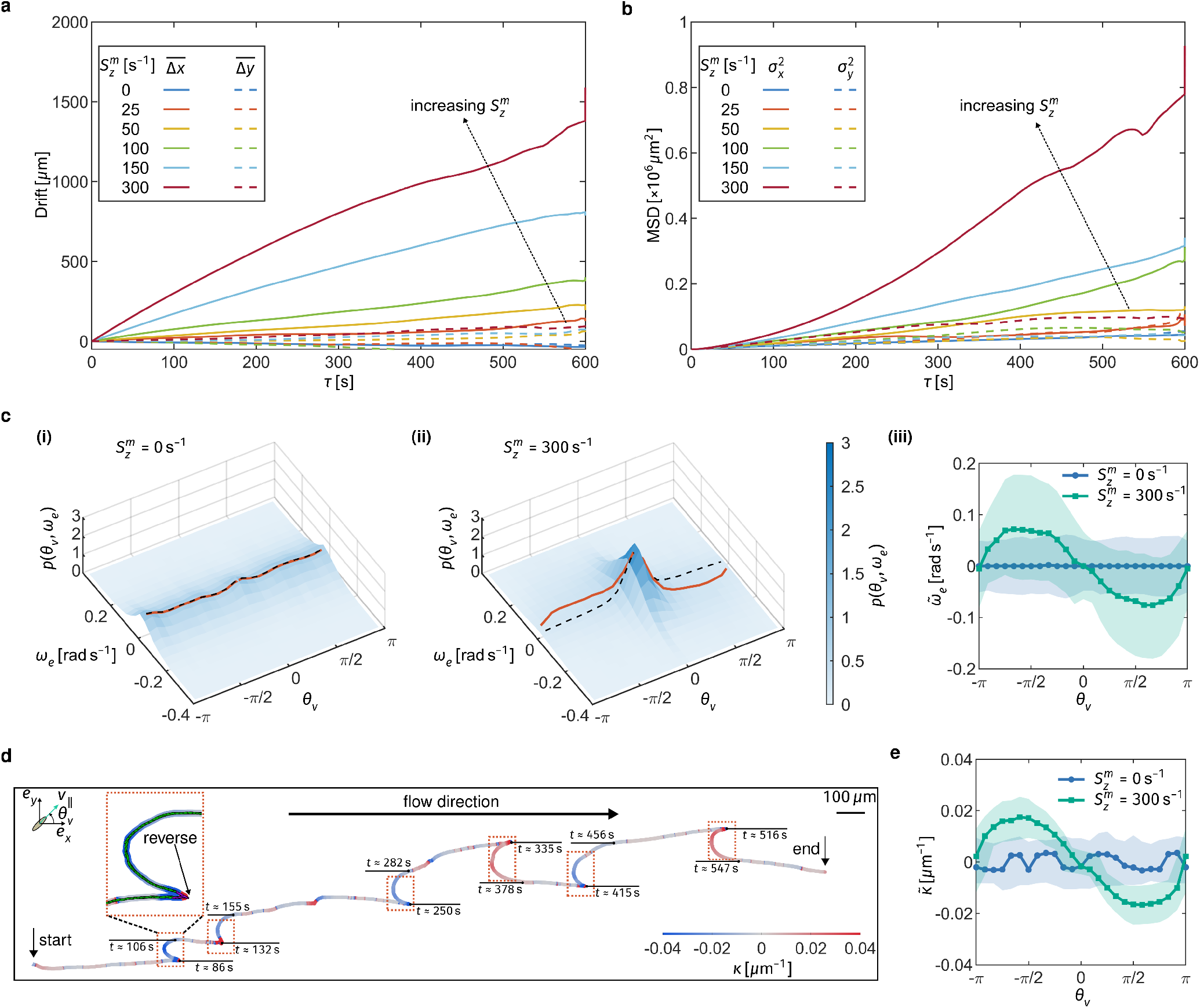
Shear-driven reorientation of gliding cells underlies downstream rheotaxis and anisotropic diffusion. **a**, Time evolution of the mean streamwise and transverse drifts 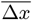 and 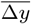, defined parallel and perpendicular to the imposed flow, respectively. Shear induces a pronounced downstream drift while leaving the mean transverse drift close to zero. **b**, Time evolution of the streamwise and transverse mean squared displacements relative to the mean drifts, 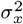 and 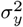. Shear strongly enhances diffusion along the flow direction. **c**, Shear-dependent angular dynamics during the GLIDE state. (**i**) and (**ii**): Joint probability density functions *p*(*θ*_*v*_, *ω*_*e*_) of angular velocity *ω*_*e*_ and gliding direction *θ*_*v*_ at 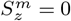 and 300 s^−1^, respectively. Black dashed lines indicate *ω*_*e*_ = 0, and red solid lines indicate the median angular velocity 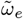 in each *θ*_*v*_ bin. (**iii**): Median angular velocity 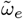 as a function of gliding direction *θ*_*v*_ . Under shear, the angular response reorients cells toward downstream gliding (*θ*_*v*_ = 0), rendering downstream motion stable and upstream motion unstable. Shaded regions indicate the interquartile range (IQR). **d**, Representative trajectory at 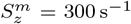, coloured by path curvature. Red dashed boxes mark REVERSE events that transiently generate upstream gliding; these are followed by rapid reorientation back toward downstream motion. Positive curvature denotes counterclockwise turning and negative curvature denotes clockwise turning. See Extended Data Fig. 7 for additional curvature-coloured trajectories at other bed shear rates. **e**, Median path curvature 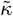 as a function of gliding direction *θ*_*v*_ . Under shear, curvature becomes strongly direction-dependent, with low curvature for downstream gliding and maximal magnitude near *θ*_*v*_ = ±*π/*2, consistent with the angular dynamics in **c**. Shaded regions indicate the IQR.

Shear flow also strongly enhanced streamwise diffusion. The streamwise mean squared displacement, 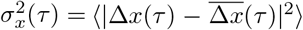, increased markedly with 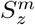, whereas the transverse mean squared displacement, 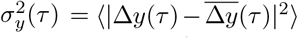, remained much smaller (Fig. 2b). Thus, shear transforms an otherwise nearly isotropic gliding process into a strongly anisotropic one. Again, removing HOP segments did not alter this conclusion (Extended Data Fig. 4).

To identify the origin of downstream rheotaxis and anisotropic diffusion, we examined the joint probability density *p*(*θ*_*v*_, *ω*_*e*_) for all GLIDE segments, where *θ*_*v*_ is the gliding direction along the cell long axis and *ω*_*e*_ is the angular velocity, both extracted from ellipsoidal tracking (Methods). At zero bed shear rate, gliding directions were nearly uniformly distributed, and the median angular velocity remained close to zero across all *θ*_*v*_ (Fig. 2c i and iii). Under shear, by contrast, gliding directions became increasingly concentrated around *θ*_*v*_ = 0, indicating downstream rheotaxis (Fig. 2c ii; Extended Data Fig. 5). This bias arose from a direction-dependent angular velocity that systematically reoriented cells towards the downstream direction, rendering downstream motion (*θ*_*v*_ = 0) stable and upstream motion (*θ*_*v*_ = ±*π*) unstable (Fig. 2c iii; Extended Data Fig. 6).

An important question is whether this angular response is governed by the gliding direction *θ*_*v*_ or by the long-axis orientation of the cell body, *θ*_*e*_ ∈ [ −*π/*2, *π/*2). These variables are not equivalent, because cells moving in opposite directions can share the same long-axis orientation. We found that, for two subpopulations gliding in opposite directions, the median angular velocity had nearly the same magnitude but opposite signs (Fig. 2c iii; Extended Data Fig. 6). Thus, near-bed shear flow acts on the instantaneous gliding direction rather than on cell orientation alone.

This mechanism is also evident in individual trajectories. A representative trajectory at 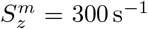 shows strong downstream migration, with streamwise displacement far exceeding transverse displacement (Fig. 2d; see Extended Data Fig. 7 for additional trajectories at other bed shear rates). Along this trajectory, the cell undergoes several REVERSE events, during which *θ*_*v*_ changes sign while *θ*_*e*_ remains continuous [31]. When a REVERSE generates upstream motion, the cell rapidly reorients back toward the downstream direction. Consequently, upstream excursions are short-lived, whereas downstream runs persist longer. Consistently, the timescale for a continuous gliding event (equivalent to the mean dwell time of the GLIDE motility state; see Extended Data Fig. 3) is comparable to the shear-induced reorientation timescale (defined as the time required to rotate from *θ*_*v*_ = 3*/*4*π* to *θ*_*v*_ = 1*/*4*π*) at intermediate to high bed shear rates (Extended Data Fig. 8), indicating a substantial downstream alignment effect.

Trajectory curvature further reveals the strong directional dependence of this reorientation. Downstream segments have low absolute curvature, whereas upstream segments are much more strongly curved (Fig. 2d; Extended Data Fig. 7). Curvature is maximal near *θ*_*v*_ ≈ ± *π/*2, consistent with the peak in 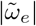 at these angles (Fig. 2c iii). The same pattern appears at the population level: 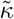 is nearly independent of *θ*_*v*_ in still water, but under shear it becomes strongly direction-dependent, with minima at *θ*_*v*_ = 0 and ±*π*, and maxima near *θ*_*v*_ = ±*π/*2 (Fig. 2e).

### 2.3 Hydrodynamic rolling converts active gliding into a downstream-restoring torque

We next sought the physical mechanism underlying the direction-dependent angular velocity and path curvature observed under shear. Two candidate explanations proved inconsistent with the data, pointing instead to a third mechanism based on shear-induced rolling during active gliding.

The first possibility is that near-bed shear flow shifts the cell–substrate contact site longitudinally towards a region of higher intrinsic raphe curvature, consistent with the common view that raphe geometry can dominate path curvature in still water for some species [31, 32]. Our observations are inconsistent with this explanation. Most importantly, it cannot explain the sign reversal of angular velocity and path curvature between cells gliding in opposite directions, because local raphe curvature alone does not determine a preferred sign of angular velocity when averaged over trajectories that include REVERSE events. In addition, the maximum path curvature expected from a typical longitudinal shift of *d*_lon_ ≈ 3.5 *µ*m (Fig. 3c ii) is around 0.01 *µ*m^−1^ (Extended Data Fig. 9a and b), notably smaller than the measured path curvature (Fig. 2e). The angular dependence is also inconsistent with this mechanism. Except for strictly upstream motion, the median longitudinal shift changes only weakly with gliding direction, whereas angular velocity and path curvature do not exhibit a comparable plateau.

**Fig. 3.**
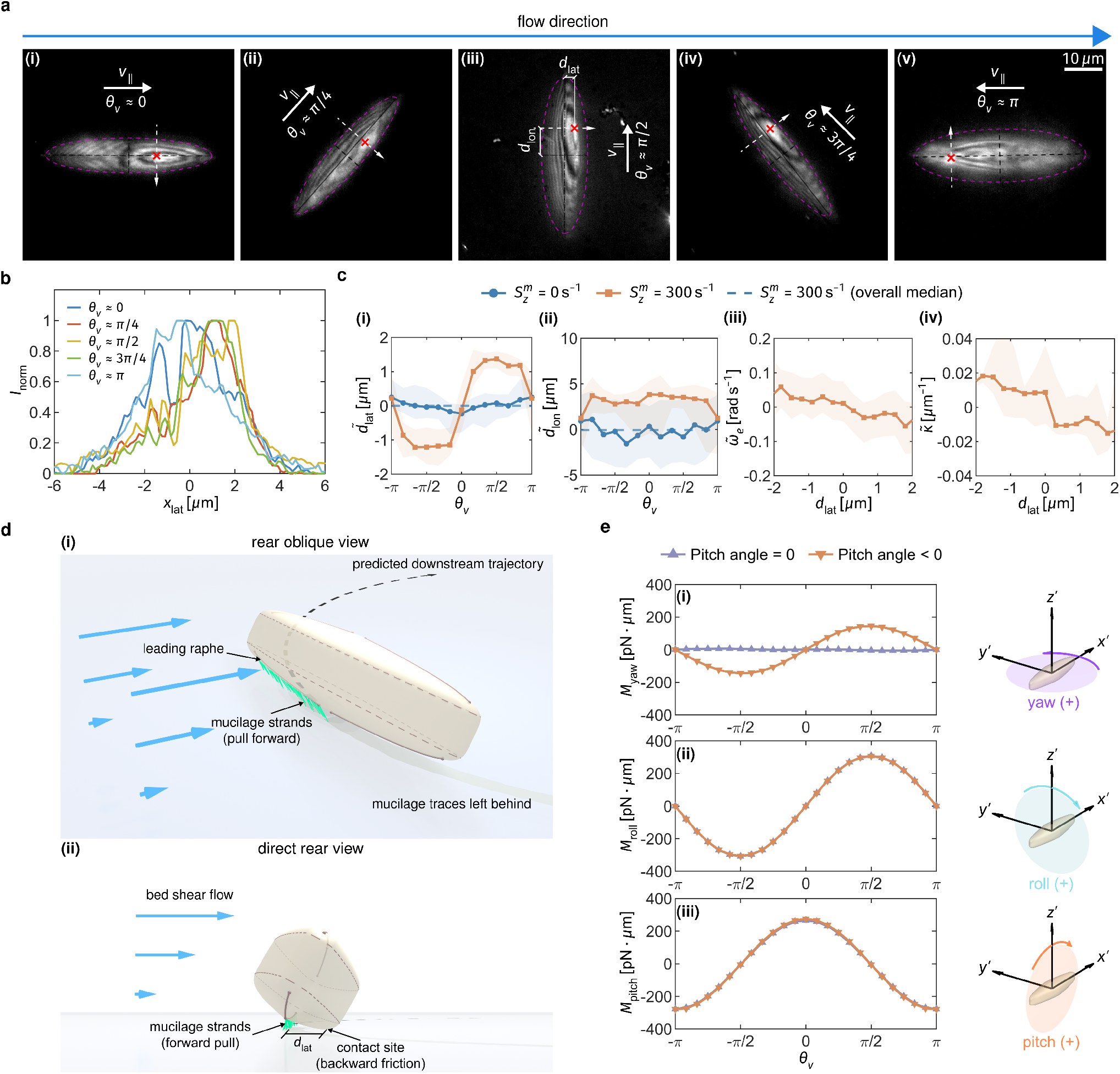
Hydrodynamic rolling shifts the contact site and converts active gliding into a downstream-restoring torque. **a**, Enhanced interference reflection microscopy images of *N. cryptocephala* gliding at different angles relative to the flow at 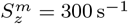. The gliding direction is indicated by the green arrow, and the frustule outline is shown in a magenta dashed line. The contact site, defined as the mean position of the five brightest pixels, is marked by a red cross. The major and minor axes are indicated by black dashed lines. The white dashed line is perpendicular to the major axis, passes through the contact site, and is oriented from left to right relative to the gliding direction. **b**, Normalised grey intensity, *I*_norm_, as a function of lateral position *x*_lat_ along the white dashed lines in **a**, with each profile normalised by its maximum intensity. Here, negative and positive *x*_lat_ indicate the left and right sides of the cell, respectively, defined with respect to the gliding direction, while *x*_lat_ = 0 corresponds to the major axis. **c**, Quantitative relation between contact site displacement and shear-induced reorientation at 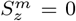 and 300 s^−1^. (**i**): Median lateral shift of the contact site, 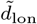, as a function of gliding direction *θ*_*v*_ . (**ii**): Median longitudinal shift of the contact site, 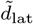, as a function of gliding direction *θ*_*v*_. (**iii**): Median angular velocity 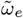 as a function of lateral shift *d*_lat_. (**iv**): Median path curvature 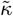as a function of lateral shift *d*_lat_. Shaded regions indicate the IQR. **d**, Schematic of the proposed mechanism. Under shear flow, hydrodynamic rolling laterally displaces the contact site during oblique gliding, so that the propulsive pull exerted through the raphe and the substrate friction are no longer collinear. This force offset generates a yaw torque that steers the cell toward downstream gliding. (**i**): rear oblique view. (**ii**): direct rear view. **e**, Hydrodynamic yaw (**i**), roll (**ii**) and pitch (**iii**) torques obtained from CFD simulations. Direct hydrodynamic yaw torque alone predicts upstream-aligning rotation, inconsistent with the measured angular response, whereas the simulated roll and pitch torques are consistent with the observed lateral and longitudinal shifts of the contact site.

The second possibility is that near-bed shear flow directly exerts a hydrodynamic yaw torque that aligns the cell downstream. To test this, we performed computational fluid dynamics (CFD) simulations for cells gliding at different in-plane directions and at either zero or negative pitch angle (Methods). For tilted configurations, the pitch angle was set to −3^°^, as estimated from the measured longitudinal shift of the contact site and z-stack imaging (Fig. 3c ii; Methods). For strictly upstream motion (*θ*_*v*_ = ±*π*), however, the pitch angle was set to zero, consistent with experiment (Fig. 3c ii). At zero pitch angle, the simulated yaw torque was negligible. At non-zero pitch angle, shear did generate a hydrodynamic yaw torque, maximal when the gliding direction was perpendicular to the flow (Fig. 3e i). Crucially, however, the torque always rotated the cell towards the upstream direction, because the lifted trailing end experienced greater fluid drag. Its sign is therefore opposite to that observed experimentally (Fig. 2c iii). This sign mismatch rules out direct hydrodynamic yaw torque as the primary origin of the observed downstream rheotaxis.

Our data instead identify a previously unrecognized mechanism: rolling induced by near-bed shear flow coupled to active gliding. For cells gliding exactly upstream or downstream, hydrodynamic symmetry predicts negligible lateral shift of the contact site (Fig. 3a i and v; Fig. 3b; Supplementary Videos 6–7). By contrast, for oblique gliding cells, near-bed shear flow produces a hydrodynamic rolling torque that laterally shifts the contact site, as revealed by IRM images and the extracted grey-value profiles (Fig. 3a ii–iv; Fig. 3b; Supplementary Video 8), and confirmed by CFD (Fig. 3e ii). The propulsive pull exerted through the mucilage acts along the raphe, whereas substrate friction is effectively applied near the contact site. In an inclined gliding posture, these two forces therefore become laterally offset. Their misalignment generates a yaw torque that rotates the cell downstream (Fig. 3d). Thus, near-bed shear flow does not reorient the cell primarily through a direct hydrodynamic yaw torque; instead, it repositions the contact site laterally so that active pulling and substrate friction form a restoring torque couple.

This interpretation is supported quantitatively by the angular dependence of the contact-site shift. The median lateral shift 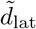 is maximal near *θ*_*v*_ ≈ ± *π/*2, where rolling torque is strongest, and approaches zero near *θ*_*v*_ = 0 and ±*π*, where hydrodynamic symmetry is restored (Fig. 3c i). Moreover, both the median angular velocity 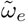 and the median path curvature 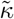 scale closely with *d*_lat_ (Fig. 3c iii and iv), indicating that the lateral offset between pull and friction is the dominant control factor shaping shear-induced reorientation.

The CFD simulations also help explain why the longitudinal shift is much larger during downstream gliding than during upstream gliding. This difference arises from the coupling between hydrodynamic pitching and the intrinsic fore–aft asymmetry of active gliding. Because propulsion and substrate adhesion are both concentrated near the leading end, the leading end acts as the main anchoring site, whereas the trailing end is more readily displaced vertically. During downstream gliding, the trailing end is lifted, producing a pronounced positive longitudinal shift 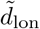 and moving the contact site closer to the leading end (Fig. 3c ii). During upstream gliding, the trailing end is instead pressed downward, resulting in a much smaller longitudinal shift. For other oblique gliding directions, the same mechanism yields a net longitudinal shift. In the absence of shear flow, no significant median longitudinal shift is detected (Fig. 3c ii; Supplementary Video 9).

Inclined gliding has also been reported in still water, possibly owing to valve-face curvature or obstacle interactions [58]. Our results show that, under near-bed shear flow, inclination is instead generated reproducibly and directionally by hydrodynamic forcing, providing the geometric asymmetry required for deterministic rheotactic reorientation.

### 2.4 A minimal stochastic model predicts shear-dependent population transport

We next asked whether a minimal stochastic model could account for the observed downstream rheotaxis and population transport (Methods). Rather than introducing direction dependence into every subprocess, we restricted it to the angular velocity and gliding speed in the GLIDE state. We also neglected the PIVOT state as an explicit model state, because it accounts for no more than 4% of all trajectory points (Extended Data Fig. 3). Instead, the net orientational randomness associated with PIVOT and other noise sources was represented by an effective reorientation angle during the REVERSE state, treated as a relaxation parameter and calibrated against the experimental orientation distribution (Methods). The HOP state was neglected entirely, as it accounts for less than 0.1% of all trajectory points (Extended Data Fig. 3).

Despite these simplifications, the model reproduced the observed shear-dependent alignment of gliding directions (Fig. 4a; Extended Data Fig. 5). At intermediate bed shear rates, the distribution of gliding directions showed a dominant downstream peak together with a smaller upstream peak due to REVERSE (Extended Data Fig. 5). As bed shear rate increased further, downstream alignment became progressively stronger, whereas upstream-directed motion became increasingly rare. The model also captured the population-level drift and diffusion at both zero and non-zero bed shear rates (Fig. 4b; Extended Data Fig. 5). In the absence of shear, cells exhibited no net drift and diffused isotropically. Under shear, they developed a mean downstream drift and anisotropic diffusion with streamwise spreading exceeding transverse spreading. Notably, this agreement was obtained without introducing any directional dependence into the transition matrix or state dwell time distributions; these were treated as isotropic, and shear entered the model predominantly through the GLIDE-state angular velocity and the direction-dependent gliding speed. This indicates that the observed downstream rheotaxis and anisotropic diffusion arise primarily from shear-induced reorientation, rather than from direction-dependent changes in state switching or persistence.

**Fig. 4.**
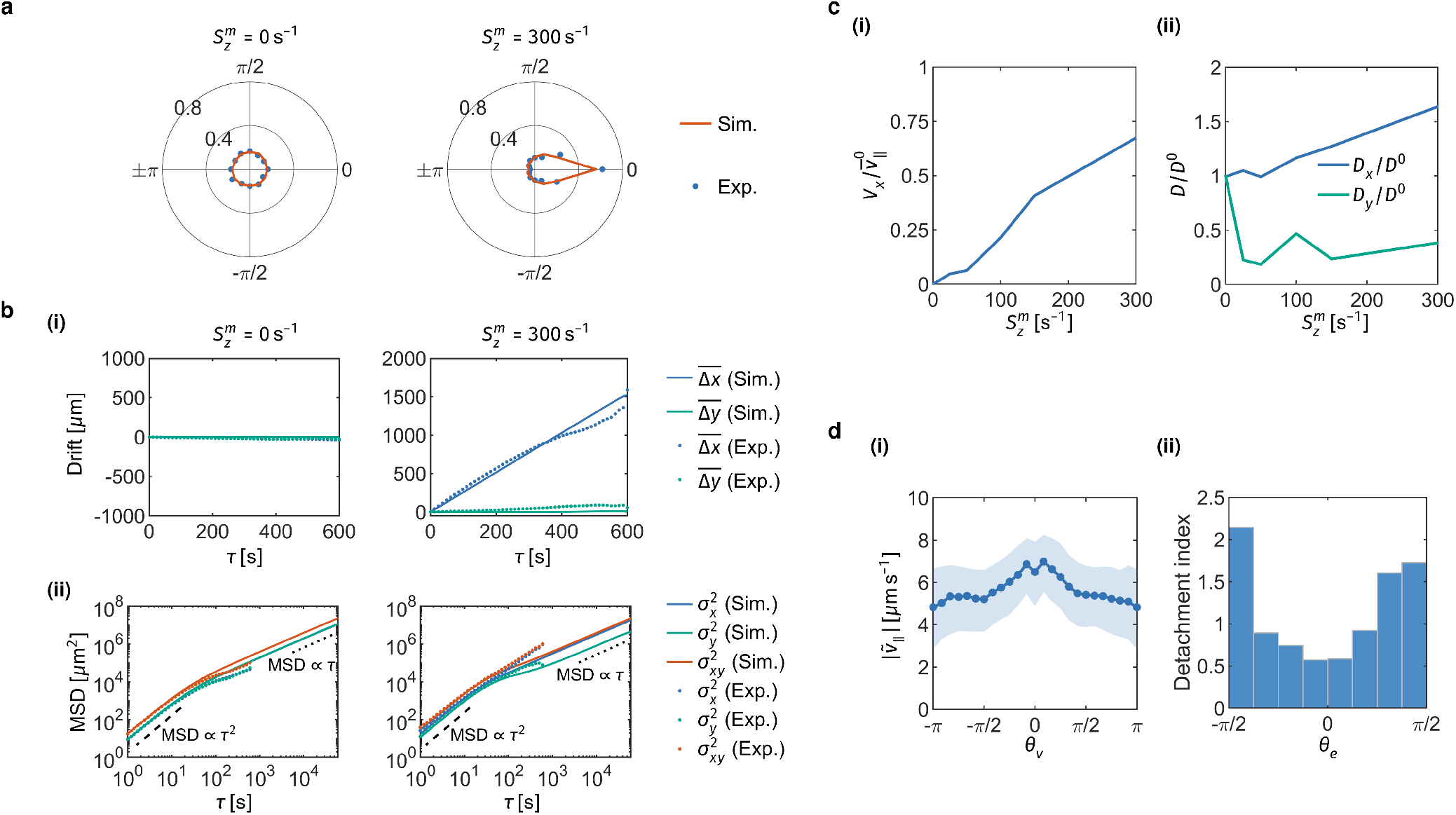
A minimal stochastic model links downstream rheotaxis to population transport. **a**, Simulated and experimental distributions of gliding direction at 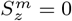 and 300 s^−1^, showing the emergence of downstream alignment under shear. **b**, Simulated and experimental displacement statistics at 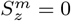 and 300 s^−1^. (**i**): Mean streamwise and transverse drifts, 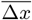 and 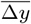, as functions of lag time, *τ* . (**ii**): Mean squared streamwise and transverse displacements relative to the mean drifts, 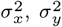, and the total mean squared displacement 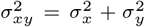. The two dashed lines in each panel indicate the ballistic scaling (MSD ∝ *τ* ^2^) and diffusive scaling (MSD ∝*τ* ). **c**, Long-time transport predicted by the stochastic model under shear flow. (**i**): Steady streamwise drift velocity, *V*_*x*_, normalised by the mean gliding speed in still water, 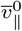, as a function of bed shear rate 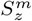. (**ii**): Steady effective streamwise and transverse diffusivities, *D*_*x*_ and *D*_*y*_, normalised by the mean diffusivity in still water, *D*^0^, as functions of bed shear rate 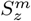. **d**, Direction-resolved motility measured experimentally. (**i**): Gliding speed as a function of gliding direction at 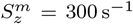; shaded regions indicate the IQR. (**ii**): Normalised detachment index as a function of long-axis orientation, *θ*_*e*_, at 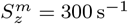. The detachment index is defined as the number of detached trajectories divided by the number of trajectory points in each *θ*_*e*_ bin, and then normalised by the corresponding value over the full *θ*_*e*_ range.

To quantify long-time transport, we next examined the lag-time dependence of the displacement statistics. At short lag times, motion was approximately ballistic, reflecting persistent gliding between stochastic state transitions. At longer lag times, repeated REVERSE events and other orientational fluctuations progressively erased directional memory, leading to a crossover to effective diffusive behaviour (Fig. 4b i and ii). Because the field of view limited the observable track length, the experimental mean squared displacements did not fully reach the asymptotic diffusive regime within the recording window. Uncertainty also increased at large lag times because the number of valid trajectory segments declined rapidly. We therefore used the validated model to extend the simulations to long times and quantify steady transport beyond the experimental window (Fig. 4c).

The effective downstream drift velocity increased with bed shear rate, reaching a maximum of approximately 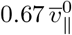 (Fig. 4c ii), where 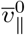 is the mean gliding speed in still water, used here as a velocity scale; the effective drift in still water is zero. This enhancement arose from two effects: shear biased cells to align downstream, and cells glided faster downstream than upstream. The speed difference approached 2 *µ*m s^−1^ at 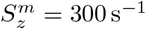 (Fig. 4d i; see Extended Data Fig. 2 for the corresponding results at other bed shear rates). Shear also made transport increasingly anisotropic. At 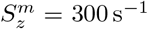, the streamwise diffusivity was enhanced by 64% relative to the mean diffusivity in still water (Fig. 4c ii). By contrast, the transverse diffusivity fell to less than half of the diffusivity in still water (Fig. 4c ii). Beyond its effect on transport, downstream alignment may also reduce the risk of scour: at 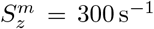, the final long-axis orientation of attached cells before detachment showed that these events occurred preferentially when the cell long axis was strongly misaligned with the streamwise direction (Fig. 4d ii).

## 3 Discussion

Our results show that benthic diatoms do not merely undergo passive perturbation or scour under shear, but can actively reorient within flow. In *Navicula cryptocephala*, near-bed shear generates downstream rheotaxis and strongly anisotropic diffusion. The underlying mechanism is mechanical: obliquely gliding cells undergo shear-induced rolling, which laterally displaces the cell–substrate contact site. This displacement makes the raphe-transmitted propulsive force non-collinear with substrate friction, thereby generating a downstream-restoring yaw torque. Thus, flow does not steer the cell primarily through a direct hydrodynamic yaw torque; instead, it reconfigures the interfacial geometry through which active gliding forces are transmitted. This new mechanism extends current descriptions of diatom gliding from a largely quiescent, cell-intrinsic framework to one in which active gliding, contact geometry and hydrodynamic forcing are explicitly coupled [31, 32, 37].

Beyond raphid diatoms, our findings identify a force-couple mechanism for directional control in surface gliding. In this mechanism, a cell is coupled to a substrate, while active propulsion and passive resistance act along spatially separated lines to generate a turning torque. In *N. cryptocephala*, this separation is generated dynamically by hydrodynamic rolling. In other gliding systems, analogous torque-generating offsets may instead arise from asymmetric adhesion, chiral surface flows, curved cell shapes, helical motor tracks or extracellular polymer transport [1, 8–10, 12, 59]. More generally, directional control during surface gliding can emerge from how active traction, passive resistance and cell geometry are spatially arranged. The resulting rheotactic direction and strength should therefore depend on cell shape, adhesion distribution, motility architecture and the susceptibility of the contact region to shear-induced rolling.

The mechanism described here is mechanically parsimonious because it does not require an explicit sensory response to flow direction. Instead, shear physically changes the force-transmission geometry through which active gliding forces act on the substrate. Such passive–active coupling provides a simple route by which surface-dwelling microorganisms can exploit near-bed flow as a directional cue. At ecological scales, near-bed shear flow shapes the architecture and spatial organization of benthic biofilms, including diatom-rich structures [33, 34, 60, 61]. Shear-induced reorientation may therefore contribute to the retention, dispersal, and anisotropic spreading of motile benthic diatoms in high-shear habitats. Together, our findings suggest that the spatial ecology of benthic microbes is shaped not only by cell-intrinsic motility mechanisms, but also by the mechanical constraints and directional cues imposed by flow near surfaces.

## 4 Methods

### 4.1 Diatom culturing

The freshwater pennate diatom *Navicula cryptocephala* (strain FACHB-2867) used in this work was obtained from the Freshwater Algae Culture Collection at the Institute of Hydrobiology (FACHB), National Aquatic Biological Resource Center, China. This strain was originally isolated from Wuhan, Hubei Province, China. Cultures were maintained in CSI medium at 25 ^°^C under a 12 h:12 h light–dark cycle with a light intensity of 2, 000 lux.

### 4.2 Microfluidic assays

Individual diatom motility under controlled bed shear rates was investigated using a custom microfluidic device. The microchannel had a serpentine geometry with three bends, a width of 2 mm, and a height of 0.1 mm. In the curved sections, the outer and inner diameters were 4 mm and 2 mm, respectively. The device was fabricated from polydimethylsiloxane (PDMS) bonded to a glass coverslip. The glass substrate was positioned at the bottom of the channel so that cells adhering to the surface could be imaged with an inverted microscope.

Fluidic connection to the microchannel was established through Luer-lock interfaces. Flow was driven by a desktop pressure source (P-PUMP02, Micro-blox Technologies) and regulated using a pressure controller (MFCS-EZ, P/N EZ-11000001, Fluigent). The imposed flow rate was monitored using a microfluidic flow meter (FLOW UNIT, model M, Fluigent).

Before each experiment, CSI medium was loaded into a reservoir bottle and connected to the entire microfluidic system, including the tubing, flow-control components, and microchannel. The system was first checked for airtightness. The pressure pump was then turned on to fill the entire system with culture medium, after which the pump was switched off. A diatom suspension was subsequently loaded into a syringe and introduced from the outlet tubing at the downstream end of the system until the cell suspension reached the observation region of the microchannel; injection was then stopped and the syringe was removed. Before loading, the diatom suspension was diluted with CSI medium so that the cell density in the observation window remained low, typically with fewer than 50 cells in the field of view in each experiment. This dilution minimised cell–cell collisions and other intercellular interactions, allowing the motility statistics to be interpreted primarily at the single-cell level. For each imposed bed shear rate, multiple independent experiments were therefore performed to obtain sufficient trajectory data, and a fresh microchannel was used between repeated experiments.

After cell loading, the device was left undisturbed for 5 min to allow cells to settle and attach to the bottom glass surface prior to flow application. The pressure pump was then turned on again to generate the flow while image acquisition was started simultaneously. During postprocessing, the first frame was defined as the first frame in which any suspended impurity particle in the field of view showed clear motion.

To generate defined shear flow, different volumetric flow rates were applied to the microchannel. The cross-sectional flow velocity profile is given by [62]

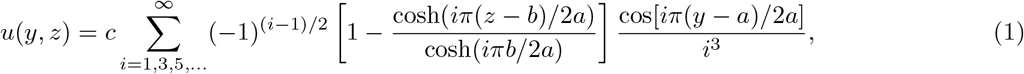

where *a* = 1 mm and *b* = 0.05 mm denote the channel half-width and half-height, respectively, and *c* is a constant determined by the prescribed volumetric flow rate. The nominal bed shear rates, 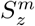, and corresponding flow rates were: (0 s^−1^, 0 nL s^−1^), (25 s^−1^, 80.7 nL s^−1^), (50 s^−1^, 161.4 nL s^−1^), (100 s^−1^, 322.8 nL s^−1^), (150 s^−1^, 484.3 nL s^−1^), and (300 s^−1^, 968.5 nL s^−1^). These controlled flow conditions enabled tracking of diatom motility on the bottom glass surface under well-defined hydrodynamic forcing.

### 4.3 Bright-field imaging for trajectory extraction and motility-state classification

Diatom motility was recorded using an inverted microscope (IX73, OLYMPUS) equipped with a 4× objective lens (UPlanFLN 4×, NA 0.13, OLYMPUS) and a custom imaging system (PCO.edge 5.5 m PIV, Excelitas PCO GmbH). During the experiments, the illumination intensity was maintained at 2, 000 lux using an LED lamp, consistent with the culturing light conditions. Imaging conditions were kept constant throughout the experiments. For each experimental condition, image sequences were acquired at 10 fps for 10 min starting from the onset of flow.

Image processing was performed in ImageJ [63]. Background fluctuations were suppressed by rolling-ball background subtraction with a ball radius of 15 pixels. Ellipsoidal particle tracking was carried out with the TrackMate plugin in ImageJ [64]. Candidate objects were detected using the Maximum Entropy thresholding detector (intensity threshold = 254). Initial morphological filtering retained objects with projected areas between 40 and 211.13 pixels2. Temporal linking was performed using the Linear Assignment Problem (LAP) tracker with a maximum frame-to-frame linking distance of 8 pixels. Gap closing was disabled to reduce false linking in dense regions. Raw trajectories were exported as CSV files and converted into physical units using a spatial calibration of 1.6 *µ*m*/*pixel and a temporal resolution of 0.1 s*/*frame. Because the cells were tracked as ellipsoidal objects, a centroid compensation step was further applied to correct small apparent centroid shifts (*<* 5 pixels) caused by orientation-dependent fitting errors.

To exclude impurities, debris, and non-motile cells, trajectories were filtered using combined morphological and kinematic criteria. Specifically, only objects satisfying the following ellipsoidal constraints were retained: major axis length *L*_*c*_ ∈ [19.2, 35.2] *µ*m (12–22 pixels), effective radius *r*_eff_ ∈ [6.4, 9.6] *µ*m (4–6 pixels), aspect ratio *AR* ∈ [1.5, 3.5], and projected area *Area* ∈ [128, 282] *µ*m^2^ ∈ (50–110 pixels2). Static impurity spots with inconsistent morphology, as well as weakly motile cells whose maximum pairwise displacement along the trajectory was smaller than 2.5*L*_*c*_ were excluded.

For motility analysis, the trajectory coordinates (*x, y*) and ellipsoidal orientation *θ*_*e*_ were processed using a segment-wise smoothing procedure to reduce noise. Trajectories were first partitioned into continuous segments based on a characteristic distance *L*_hop_ for the detection of a HOP event. The orientation angle *θ*_*e*_ was unwrapped and smoothed using a first-order Savitzky–Golay filter. For the spatial coordinates, segments were smoothed using a 5-frame moving-average filter three times. Translational velocity **v** = (*v*_*x*_, *v*_*y*_) and angular velocity *ω*_*e*_ were computed using central finite-difference schemes with step sizes of 10 and 2 frames, respectively, with appropriate adjustments near segment boundaries.

Motility was classified into five states using a hierarchical algorithm adapted from ref. [31]: GLIDE (G), STOP (S), REVERSE (R+ and R−), PIVOT (P), and HOP (H), with the latter defined in this study. The classification was primarily based on the normalised parallel velocity *v*∥ _,norm_, defined as the velocity component along the major axis normalised by the mean gliding speed.

- **GLIDE and STOP**. A sliding time window (4 s) was used to evaluate the sign stability of *v*∥ _,norm_. A segment was classified as GLIDE when the sign remained unchanged and the mean *v*∥ _,norm_ exceeded 0.025; otherwise it was classified as STOP.
- **REVERSE**. A REVERSE event was identified by a sign change in *v*∥ _,norm_ within a continuous segment. Segments with repeated oscillations were further partitioned into discrete directional intervals using a sign-flip index rule.
- **PIVOT**. Rotational transitions were identified from the pole-velocity ratio,

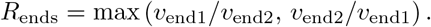

A segment was classified as PIVOT when *R*_ends_ *>* 1.5 for at least three consecutive frames, indicating substantial rotation with minimal centre-of-mass translation.
- **HOP**. Discontinuous jumps with displacement greater than *L*_hop_ = 15 *µ*m were classified as HOP.

Across all bed shear rate conditions, the numbers of retained trajectories at 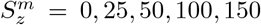, and 300 s^−1^ were 234, 157, 250, 225, 307, and 464, respectively. The corresponding mean trajectory durations were 245.0, 268.3, 230.2, 249.3, 241.9, and 178.0 s.

### 4.4 Interference reflection microscopy

Interference reflection microscopy (IRM) was used to image the contact region between diatom cells and the glass substrate. Imaging was performed on an inverted fluorescence microscope (Ti-2, Nikon). A high-numerical-aperture total internal reflection fluorescence (TIRF) objective (Apo TIRF 60× oil, NA 1.49, Nikon) was used. Non-polarized IRM images were acquired using a narrow-band interference filter, such that image contrast arose from the interference of reflected light at the sample–glass interface. Illumination was provided by a mercury lamp (INTENSILIGHT C-HGFIE, Nikon) at 532 nm. The aperture diaphragm was set to 75% open. All optical parameters were kept fixed throughout data acquisition to ensure consistency between measurements. Image sequences were acquired using a sCMOS camera (ORCA-Flash4.0, Hamamatsu) with an exposure time of 200 ms and a frame rate of 5 fps.

Time-lapse IRM image sequences were processed using a custom workflow to extract diatom body geometry, cell–substrate contact bright spots, and cell trajectories. Raw Nikon ND2 files were first converted into single-frame TIFF images using Bio-Formats in Fiji, with each time point exported as an individual image. To identify the diatom frustule in IRM images, a pixel-classification model was trained in Fiji using the Trainable Weka Segmentation 2D plugin [65]. Representative frames were manually annotated to distinguish frustule pixels from background, and the trained classifier was then applied to all TIFF frames to generate frustule probability maps. The resulting probability images were converted into binary masks by thresholding, followed by particle analysis to isolate individual diatom outlines. For each segmented cell, an ellipse was fitted to the binary object to obtain the cell centre coordinates, major and minor axis lengths, and ellipse orientation angle for each frame. This representation is consistent with the bright-field tracking workflow, in which cell position and orientation were likewise described by fitted ellipse parameters.

For each identified diatom, frame-resolved segmentation contours were exported as regions of interest (ROIs) sets and linked to the original IRM images. Bright-spot extraction was then performed on the original, unprocessed IRM frames. For each frame, the frustule ROI was used as a mask, and pixel intensities within the ROI were ranked in descending order. The mean position of the brightest five pixels was used as the representative contact site. The fitted ellipse parameters and the representative contact site were then paired by frame number to reconstruct the trajectory and corresponding contact site dynamics of each individual cell. Frames with missing detections were excluded, and discontinuities in frame number were used to split trajectories into independent continuous segments.

To relate the representative contact site to cell geometry, the displacement vector from the ellipse centre to the bright-spot position was projected onto the major and minor axes of the fitted ellipse. The major-axis direction was oriented to be consistent with the instantaneous direction of motion, and the resulting projections were used to define the longitudinal and lateral offsets of the contact site relative to the cell body. In addition, local trajectory curvature was estimated from the centre trajectory using the same sliding-window circle-fitting procedure as in the bright-field experiments. For each trajectory, only points with speeds greater than 20% of the median speed across all trajectory points were retained for analysis. This yielded 13,160 effective trajectory points for 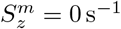 and 2,243 for 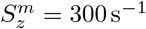.

### 4.5 Path and raphe curvature assessment

To decouple path tortuosity from fluctuations in translational speed, we estimated the local trajectory curvature *κ* using a spatial-window method. For each trajectory point *j*, we first computed the cumulative arc length 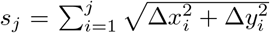, and then defined a local neighbourhood by the arc-length window [*s*_*j*_ −0.5*d*_curv_, *s*_*j*_ +0.5*d*_curv_], where *d*_curv_ is the distance window size.

A circle was fitted to all coordinates within this window by a linear least-squares minimisation of the form:

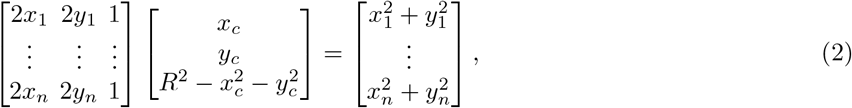

where (*x*_*c*_, *y*_*c*_) is the fitted circle centre and *R* is the radius of curvature. The local curvature magnitude was then calculated as

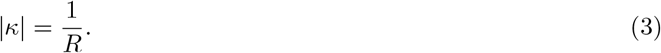

The sign of *κ* was assigned according to the turning direction of the trajectory, with positive values denoting counterclockwise curvature and negative values denoting clockwise curvature.

Because the fitting window was defined in arc length rather than in time, this procedure provides a purely geometric measure of path tortuosity and remains robust even when the diatoms move with strongly non-uniform speeds. The distance window size was set to *d*_curv_ = 30 *µ*m for both bright-field and IRM trajectory analysis.

For raphe curvature estimation, more than 400 points were manually digitized along one raphe branch on one side of each cell. The extracted raphe contour was then smoothed and resampled using a spline-based procedure to obtain a uniform point density of 20 points per *µ*m of arc length. Raphe curvature was subsequently calculated using the same circle-fitting method as for path curvature, with an arc-length window of 4 *µ*m.

### 4.6 Scanning electron microscopy

Diatom samples were pretreated before scanning electron microscopy (SEM) observation to remove non-diatom impurities, other microorganisms, and intracellular organic materials, thereby obtaining clean siliceous frustules for morphological examination. In this study, frustule cleaning was carried out using microwave digestion.

Briefly, the diatom suspension was first mixed thoroughly by gentle shaking, and 10 mL of the suspension was transferred to a centrifuge tube. The sample was centrifuged at 1, 500 rpm for 8 min, the supernatant was discarded, and the pellet was transferred to a digestion tube. Then, 10 mL of concentrated nitric acid was added to the digestion tube. After the tube was tightly sealed, it was placed in a microwave digestion system and digested at 180 ^°^C for 2 h. After digestion, the tube was opened in a fume hood and allowed to cool to room temperature. The digested sample was then transferred to a centrifuge tube and centrifuged again at 1, 500 rpm for 8 min. After removing the supernatant, distilled water was added to a final volume of 10 mL, and the sample was thoroughly resuspended. This washing step was repeated 5–6 times until the supernatant was approximately neutral.

After the final centrifugation, the supernatant was removed using a 1 mL pipette. Subsequently, 0.5 mL of absolute ethanol was added, and the pellet was resuspended thoroughly. The cleaned sample was then transferred to a 1.5 mL microcentrifuge tube. The original centrifuge tube was rinsed with absolute ethanol to recover any remaining material, and all rinses were combined in the same microcentrifuge tube for storage until SEM preparation.

An aliquot of the cleaned suspension was dropped onto an SEM stub and dried under an infrared lamp. The dried samples were then sputter-coated with gold to improve surface conductivity prior to imaging. SEM images were acquired using a scanning electron microscope (SU8010, Hitachi) operated at an accelerating voltage of 2 kV and a working distance of 7.6 mm.

### 4.7 Z-stack imaging

Three-dimensional z-stack scanning was performed to acquire volumetric image stacks of the sample. Z-stack data (axial step size 93.75 nm) were acquired using a confocal microscope (A1R HD25, Nikon) equipped with a 100× silicone-oil immersion objective (CFI SR HP Plan Apo Lambda S 100XC Sil, Nikon). Excitation was provided at 561 nm.

For each measurement, sequential optical sections were collected along the *z* direction and assembled into a three-dimensional image stack for subsequent analysis. The three-dimensional diatom geometry used in the CFD simulations was reconstructed from these confocal z-stacks. We first identified the optical section with the largest in-plane area, which is approximately parallel to the valve surfaces and represents the maximum projected cross-section of the frustule. To derive an idealised and representative diatom frustule, the maximum-area section was symmetrised along the major and minor axes and further subjected to centrosymmetry, yielding a reference cross-sectional contour (Extended Data Fig. 9e). The full three-dimensional geometry was then reconstructed by scaling this reference contour symmetrically toward the upper and lower sides according to the area ratio between each z-stack layer and the maximum-area layer (Extended Data Fig. 9f). This procedure yielded a smoothed, idealised diatom geometry for CFD simulations (Extended Data Fig. 9c and d). This reconstructed diatom geometry also allowed us to determine the non-zero pitch angle corresponding to the longitudinal shift of the contact site measured by IRM.

### 4.8 Computational fluid dynamics

Three-dimensional flow simulations were performed using the open-source solver PetIBM [66], which is designed for incompressible-flow simulations with an immersed-boundary treatment of complex solid geometries. In brief, the solver advances the incompressible Navier–Stokes equations on a Cartesian grid and imposes the no-slip condition on the immersed body through a Lagrangian representation of the solid surface coupled to the Eulerian flow field. In the present study, this framework was used to resolve the near-body flow around a gliding diatom under an imposed representative shear rate of 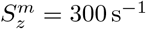.

The computational domain size was set to (6*L*_*c*_ × 6*L*_*c*_ × 5*H*_*c*_) in the streamwise, spanwise, and wall-normal directions, respectively, where the characteristic diatom length was *L*_*c*_ = 32 *µ*m and the characteristic diatom height was *H*_*c*_ = 6.1 *µ*m. This corresponds to a physical domain of 192 *µ*m×192 *µ*m×30.5 *µ*m. Grid refinement was applied in the vicinity of the diatom: in the streamwise and spanwise directions, the region extending to ±1.5*L*_*c*_ around the cell centre was uniformly refined with a grid spacing of 0.0075*L*_*c*_, i.e. 0.24 *µ*m. To avoid numerical instability associated with direct contact between the diatom and the bottom wall, the minimum wall-normal gap between the lowest point of the diatom and the substrate was set to 0.1 *µ*m, and this gap was resolved using five equally spaced grid cells. Within a wall-normal distance of 2*H*_*c*_, the grid spacing was approximately 0.20 *µ*m; from 2*H*_*c*_ to the upper boundary, the grid spacing was approximately 0.60 *µ*m. The total number of grid cells was 2.375 ×10^7^.

To avoid excessively stiff immersed-boundary forcing while maintaining geometric fidelity, two Lagrangian points were placed in each Eulerian cell intersected by the shell surface, resulting in a representation of the diatom surface with 13,236 Lagrangian points. The lower and upper boundaries were prescribed as Dirichlet velocity boundaries. In the diatom-fixed frame of reference, the lower boundary velocity was set opposite to the measured gliding velocity of the diatom, whereas the upper boundary velocity was prescribed as the sum of the opposite gliding velocity and the background shear contribution, taken as 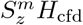, where 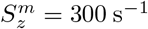 is the representative substrate shear rate and *H*_cfd_ = 5*L*_*c*_ is the computational domain height. This assumes that the imposed shear rate remains approximately constant over the computational height, which is a reasonable approximation for the present near-wall configuration. The two spanwise boundaries and the two streamwise boundaries were treated as periodic boundaries. Under this setup, the imposed background flow corresponded to a Couette-like shear generated by the relative velocity difference between the lower and upper boundaries.

Two groups of simulations were carried out. In the first group, the diatom pitch angle was set to 0^°^. In the second group, the pitch angle was set to −3^°^, except for the case of *θ*_*v*_ = *π*, for which a pitch angle of 0^°^ was used, consistent with the IRM observations (Fig. 3c ii). The imposed gliding speed depended on the gliding direction and was prescribed from the Laplace fits to the experimental velocity distributions extracted from cell trajectories. Using the characteristic scales of the diatom 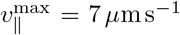 and *L*_*c*_ = 32 *µ*m, together with the kinematic viscosity of water *ν* = 10^−6^ m^2^ s^−1^, the Reynolds number is estimated as 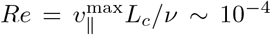, indicating a creeping-flow regime in which inertial effects are negligible. Owing to the extremely small Reynolds number, the flow converged rapidly to a steady state. The total simulated physical time was approximately 2.1 × 10^−3^ s, corresponding to 3, 000 time steps. The hydrodynamic yaw, roll, and pitch torques were computed about the diatom centroid and evaluated from the time-averaged values over the final 5 time steps after the solution had reached steady state.

### 4.9 Individual-based stochastic model

To examine whether the observed downstream rheotaxis and anisotropic diffusion can be reproduced from a minimal set of experimentally calibrated motility rules, we developed an individual-based stochastic model for surface-associated diatom migration. The model combines stochastic switching between discrete motility states with continuous updates of cell position and orientation, in a state-based stochastic framework adapted from previous models for gliding diatoms [31] and extended here to account for shear-induced downstream rheotaxis. Each simulated cell is represented by its planar position (*x*(*t*), *y*(*t*)), planar body axis orientation *θ*(*t*) ∈ [ −*π, π*), and a polarity variable *s*(*t*) ∈ {+1, −1} (with the initial gliding direction assigned as positive polarity). The instantaneous gliding direction *θ*_*v*_ was therefore defined as *θ* for *s* = +1 and *θ* +*π* for *s* = −1. This polarity variable is necessary because raphid diatoms have no morphologically fixed front or back, so REVERSE can change the direction of translation while leaving the body axis itself continuous.

The full behavioural repertoire identified experimentally consists of four stereotyped states, GLIDE, REVERSE, STOP and PIVOT (consistent with previous state-based analyses of diatom motility [31]), together with a fifth state unique to shear flow, HOP. In the present model, however, we explicitly retain only three effective states: GLIDE, REVERSE, and STOP. We neglect the HOP state, as it contributes less than 0.1% of all classified trajectory points in our data set (Extended Data Fig. 3). The PIVOT state was also omitted as an explicit state because it accounted for no more than 4% of all classified trajectory points (Extended Data Fig. 3). Instead, the effect of PIVOT was coarse-grained into an effective angular reorientation applied during the REVERSE state, representing the combined contribution of PIVOT-associated turning and other unresolved rotational fluctuations. Successive motility episodes were generated by stochastic transitions between states using an experimentally calibrated transition matrix. Specifically, after completion of each episode, the next state was sampled from a stochastic transition matrix *P* renormalised after exclusion of the PIVOT state, whose entries give the conditional probabilities of transitioning from the current state to the next one. In our implementation, all simulated trajectories were initialised in the GLIDE state, with the initial long-axis orientation *θ*_0_ distributed uniformly over [ −*π, π*).

The duration of each motility episode was drawn from a state-dependent dwell time distribution fitted to the experimental data. For the GLIDE and STOP states, dwell times were sampled from exponential distributions. For the REVERSE state, the episode duration was sampled from a Laplace distribution truncated to positive values (Extended Data Fig. 3). The model was integrated with a fixed time step Δ*t* = 0.1 s.

During the GLIDE state, the cell translated strictly along its major axis, consistent with previous modelling assumptions that tangential motion dominates and that translational dynamics are aligned with the long cell axis [31]. The position was updated as follows:

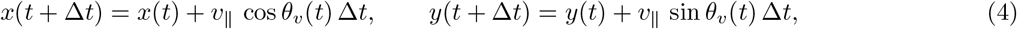

where *v >* 0 is the gliding speed. In the simulations, a single speed was sampled for each GLIDE episode and then held constant during that episode. To account for the experimentally observed dependence of gliding speed on the instantaneous gliding direction, *v*_∥_ was drawn from a Laplace-type function fitted to the data:

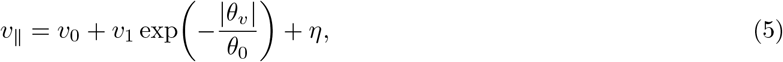

where *v*_0_, *v*_1_, and *θ*_0_ are fitted parameters, and 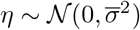 is a Gaussian random term (Extended Data Fig. 2). Negative sampled speeds were rejected and resampled to ensure *v >* 0.

Another key ingredient of the model was introduced through the angular dynamics in the GLIDE state. Rather than imposing orientation dependence on all subprocesses, we assumed that the dominant effect of shear flow enters through a deterministic angular velocity depending on the instantaneous gliding direction,

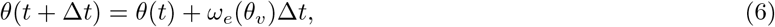

with

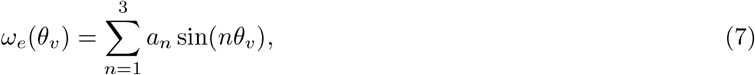

where the coefficients *a*_*n*_ were fitted independently for each bed shear rate (Extended Data Fig. 6). This Fourier representation captures the experimentally measured dependence of angular velocity on gliding direction and provides a minimal description of the downstream-restoring reorientation observed under shear. In still water, the fitted coefficients were set to zero so that no directional bias was imposed.

The REVERSE state was implemented as a finite-time polarity-switching event. Its total duration was sampled as described above, and the polarity *s* was flipped halfway through the episode. At the instant of polarity inversion, the cell orientation was additionally perturbed by a small angular offset Δ*θ*, introduced as an effective relaxation parameter. Under shear, the sign of this offset was chosen so that the post-REVERSE gliding direction was slightly shifted toward the downstream direction, consistent with the experimentally observed net reorientation across REVERSE events; at zero shear, the sign was assigned randomly with equal probability. This term provides a coarse-grained representation of the weak net turning associated with neglected PIVOT-like events and other unresolved angular noise during REVERSE.

This implementation implies a weak downstream bias during REVERSE under shear, but this bias was data-motivated. Representative trajectories (Extended Data Fig. 7) and population-level statistics of orientation change across REVERSE events (Extended Data Fig. 10a) both support such a tendency. The calibrated offset remained small, with Δ*θ* = 1^°^ for 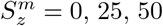, and 100 s^−1^, and Δ*θ* = 5^°^ for 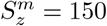 and 300 s^−1^. Sensitivity analysis further showed that a model with GLIDE-state angular bias only already reproduced the measured gliding-direction distribution and downstream drift closely, whereas a model with REVERSE-state bias only differed markedly from both the full model and experiment (Extended Data Fig. 10b). Thus, the dominant origin of rheotaxis in the stochastic model is the downstream-restoring angular velocity during GLIDE fitted to experimental data, while the REVERSE-associated offset provides only a secondary correction, mainly reducing the persistence of the upstream-oriented peak created by polarity switching.

The STOP state was modelled as a pause with no translation or rotation. For each bed shear rate, *N* = 2, 000 independent trajectories were simulated in parallel. Simulations were carried out for either 600 s, for direct comparison with experimental measurements, or 6 ×10^4^ s, to quantify long-time transport. Based on the simulated trajectories, we calculated the mean drift displacement and the mean squared displacement about the drift. For the long-time transport analysis, linear fits over the interval *τ* ∈ [3.3 ×10^4^ s, 4.3 ×10^4^ s] were used to extract the asymptotic streamwise drift velocity and diffusivity.

The model was intentionally kept minimal: shear dependence entered only through the angular dynamics during GLIDE fitted to experimental data and a weak effective reorientation during REVERSE. Despite this simplification, it reproduced the measured gliding-direction distributions, downstream drift, and anisotropic diffusion across bed shear rates.

## Acknowledgements

We acknowledge helpful discussions with Prof. K.Y. Wan and Prof. Q.X. Liu. We thank Prof. P. Yu and Y.B. Tian for assistance with SEM imaging, and J.Y. Wang for assistance with IRM and z-stack imaging. B.W. also acknowledges the hospitality of the Department of Applied Mathematics and Theoretical Physics, University of Cambridge, during his visit.

## Data availability

The data that support the findings of this study will be made available upon the acceptance of this manuscript.

## Code availability

All code used for analysis will be made available upon the acceptance of this manuscript.

## Author contributions

All authors contributed substantially to the work presented in this manuscript. B.W., L.Z. and J.W. conceived the project. B.W., S.D. and L.Z. carried out the experiments. B.W., S.D. and L.Z. analysed the experimental results. B.W., W.J., X.G. and R.H. performed the numerical simulations. T.J.P. contributed to the discussion and interpretation of key results. L.Z. and J.W. supervised the study. B.W. and L.Z. wrote the manuscript. All authors reviewed the manuscript.

## Extended Data Figures

**Extended Data Fig. 1.**
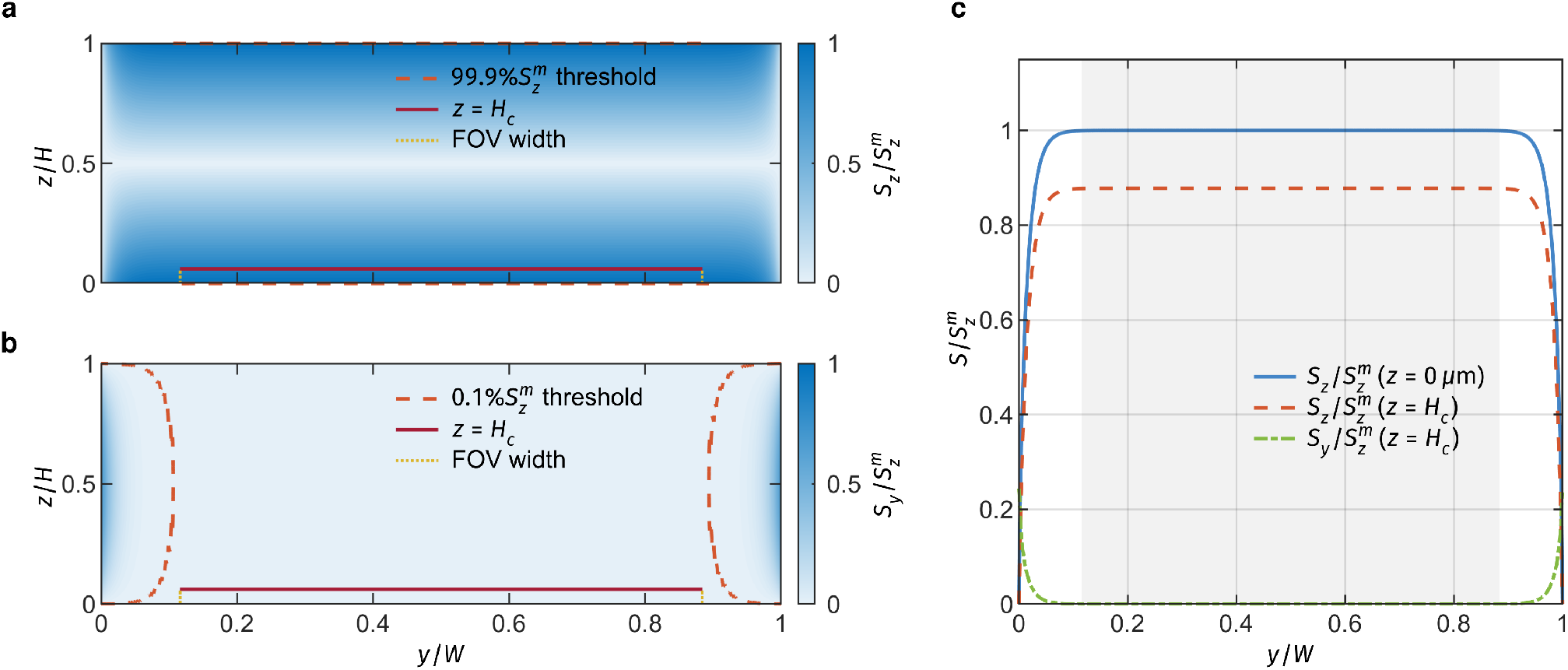
Distribution of normalised vertical and transverse shear rates across the channel cross-section. **a**, Contour map of the normalised vertical shear rate 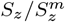. The red dashed line represents the 99.9% contour of the maximum bed shear rate 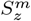. The red solid line indicates the height *z* = *H*_*c*_, where *H*_*c*_ = 6.1 *µ*m corresponds to a typical cell height. The vertical dotted lines delimit the width of the field of view (FOV). **b**, Contour map of the normalised transverse shear rate 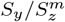. The red dashed line denotes the 0.1% level relative to 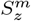. **c**, Comparison of shear rate profiles across the channel width. The plot shows the vertical shear rate at the bed (*z* = 0 *µ*m) and at *z* = *H*_*c*_, together with the transverse shear rate at *z* = *H*_*c*_. The grey shaded region indicates the width of the FOV.

**Extended Data Fig. 2.**
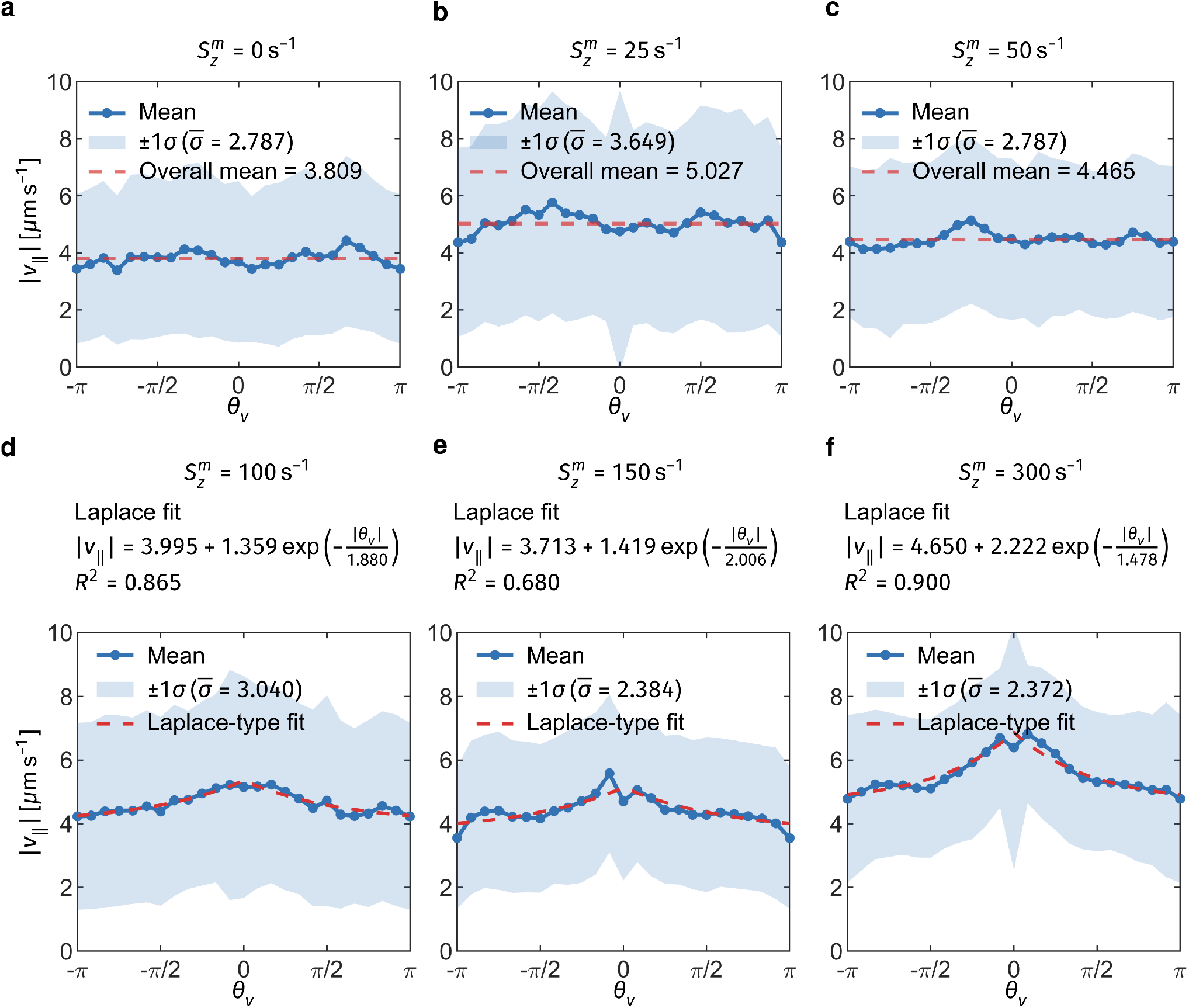
Dependence of the gliding speed |*v*_∥_| on gliding direction *θ*_*v*_ under different bed shear rates. For 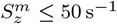, the gliding speed is approximated by a direction-independent mean value, whereas for 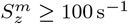, its directional dependence is represented by a Laplace-type function. **a**–**f**, 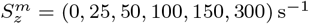.

**Extended Data Fig. 3.**
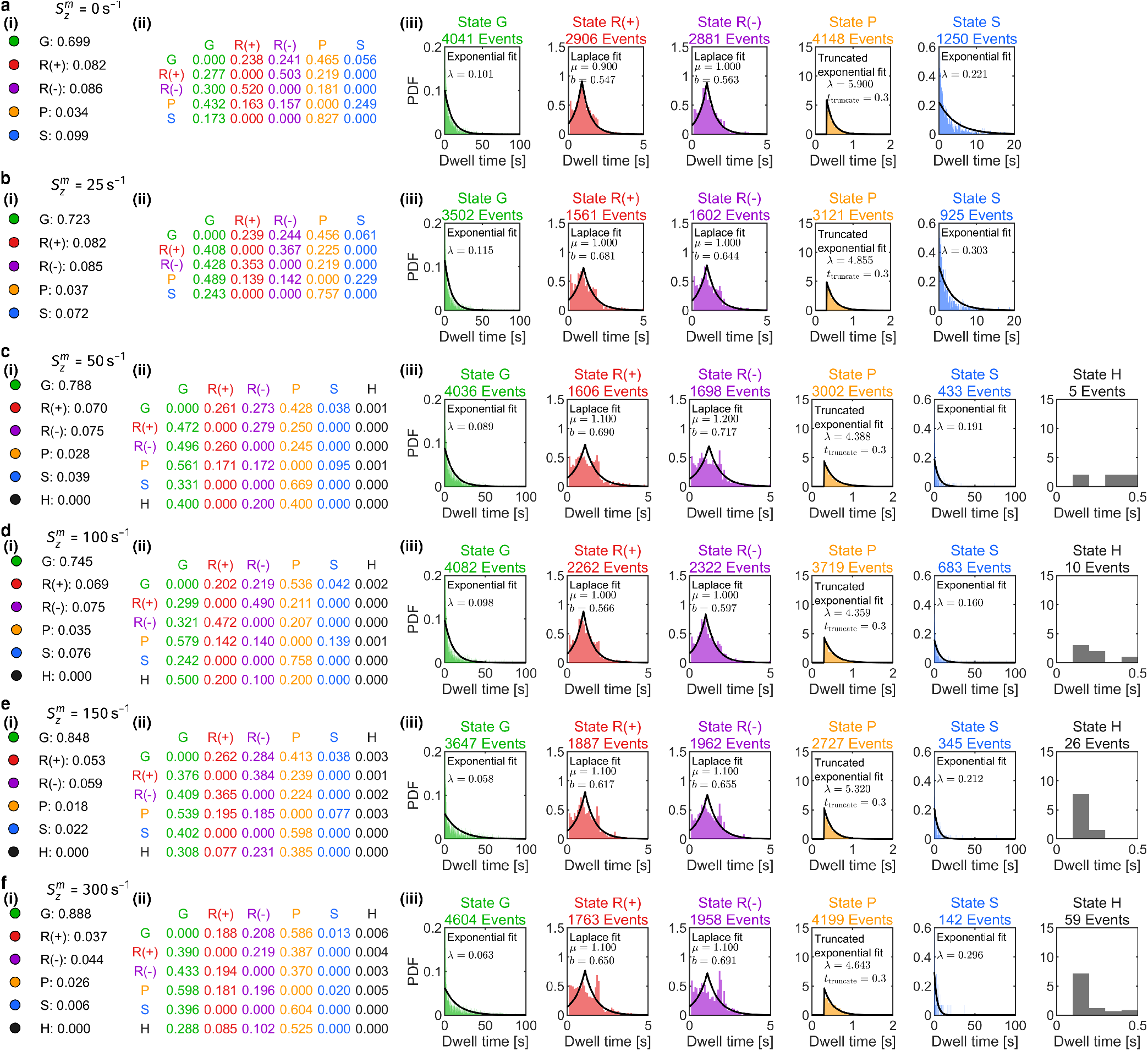
Motility-state statistics under different bed shear rates. Columns **(i)**–**(iii)** show the fraction of total trajectory points assigned to each motility state, the Markov transition matrix, and the dwell time distributions for all motility states, respectively. **a**–**f**, 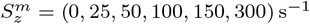.

**Extended Data Fig. 4.**
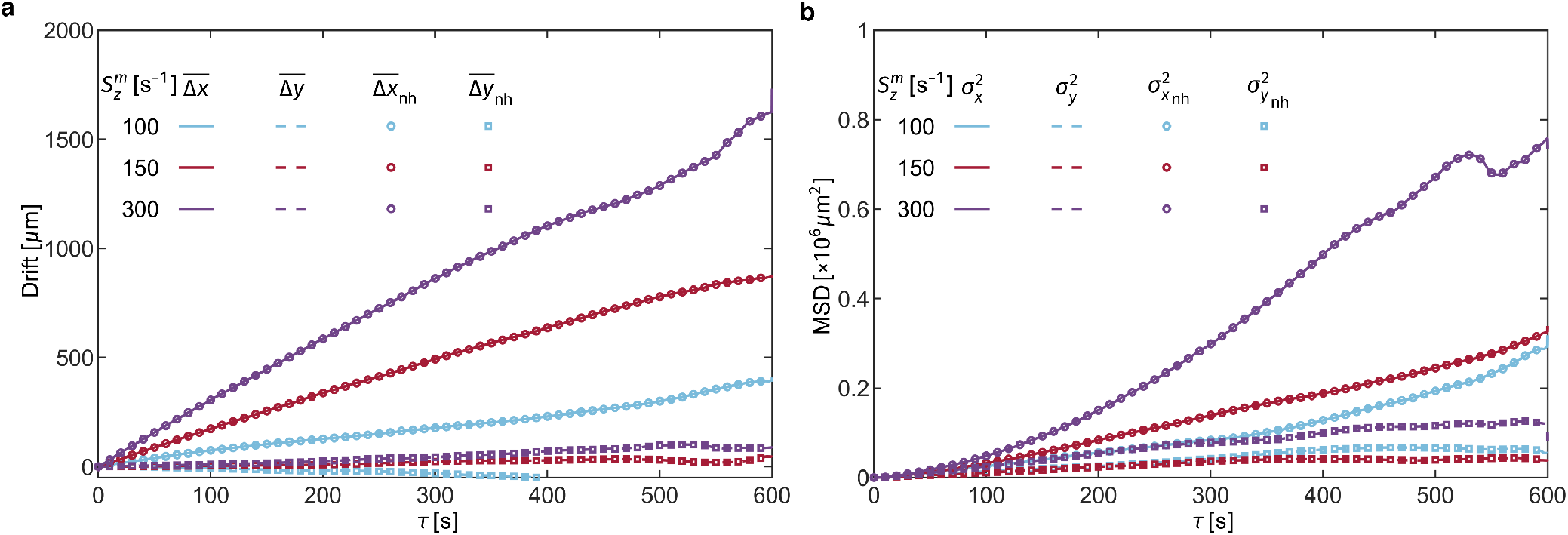
Sensitivity of downstream rheotaxis and anisotropic diffusion to HOP events. **a**, Time evolution of the mean streamwise and transverse drifts, 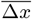 and 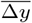, for the full dataset and for the subset excluding all trajectory segments containing HOP events (subscript ‘nh’). **b**, Mean squared streamwise and transverse displacements relative to the mean drifts, 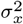 and 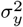, for the full trajectories and for the non-hopping trajectory segments, 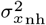 and 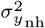 .

**Extended Data Fig. 5.**
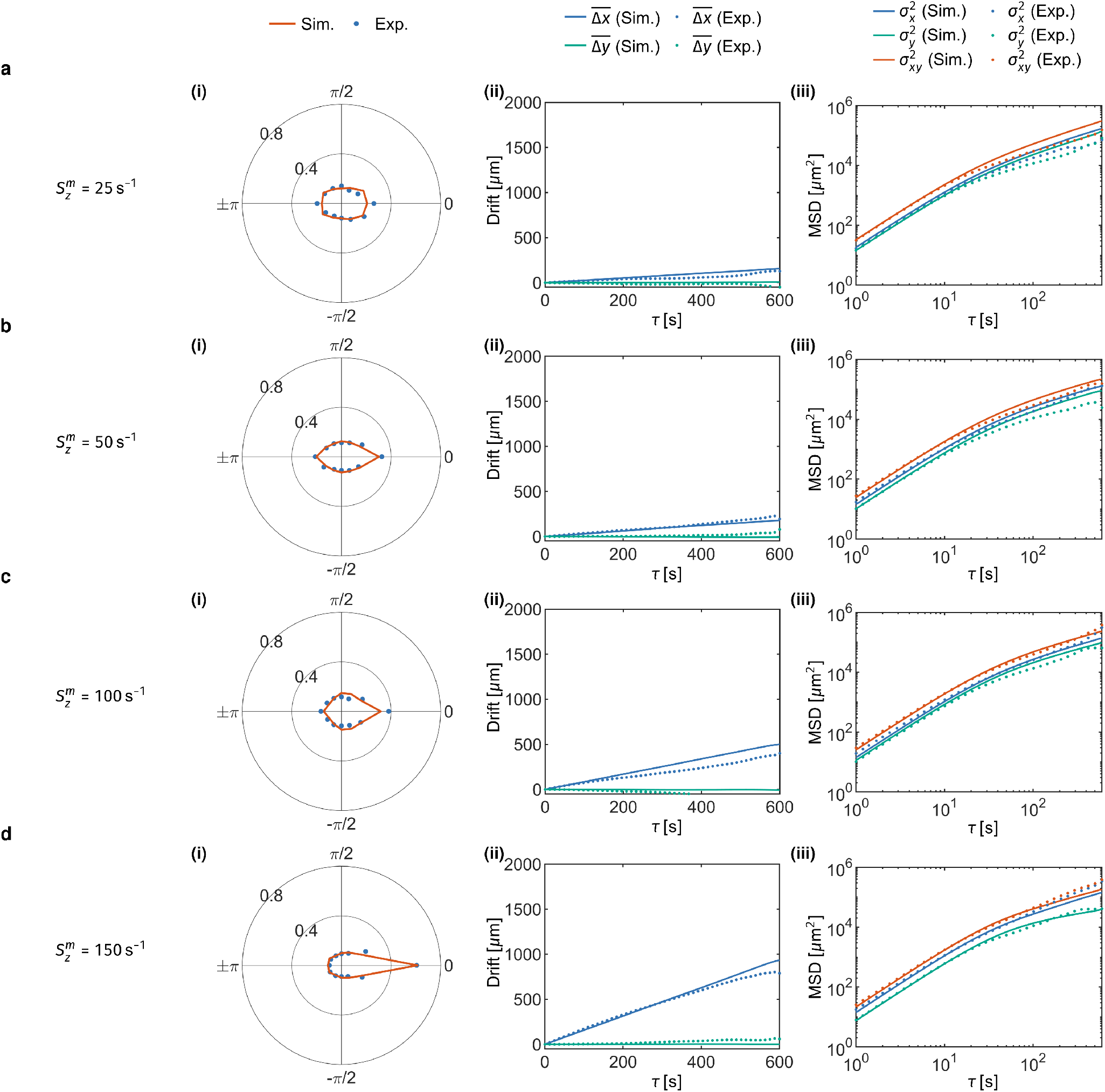
Validation of the stochastic model for other bed shear rates. Columns **(i)**–**(iii)** show the simulated and experimental results. Column **(i)** shows the distribution of the gliding direction, *θ*_*v*_ . Column **(ii)** shows the mean streamwise and transverse drift displacements, 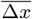 and 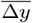. Column **(iii)** shows the mean squared streamwise and transverse displacements relative to the mean drift displacements, 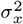 and 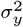, as well as the total mean squared displacement, 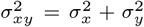. **a**–**d**, 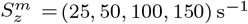 . The legends are the same as those in Fig. 4**a** and **b**.

**Extended Data Fig. 6.**
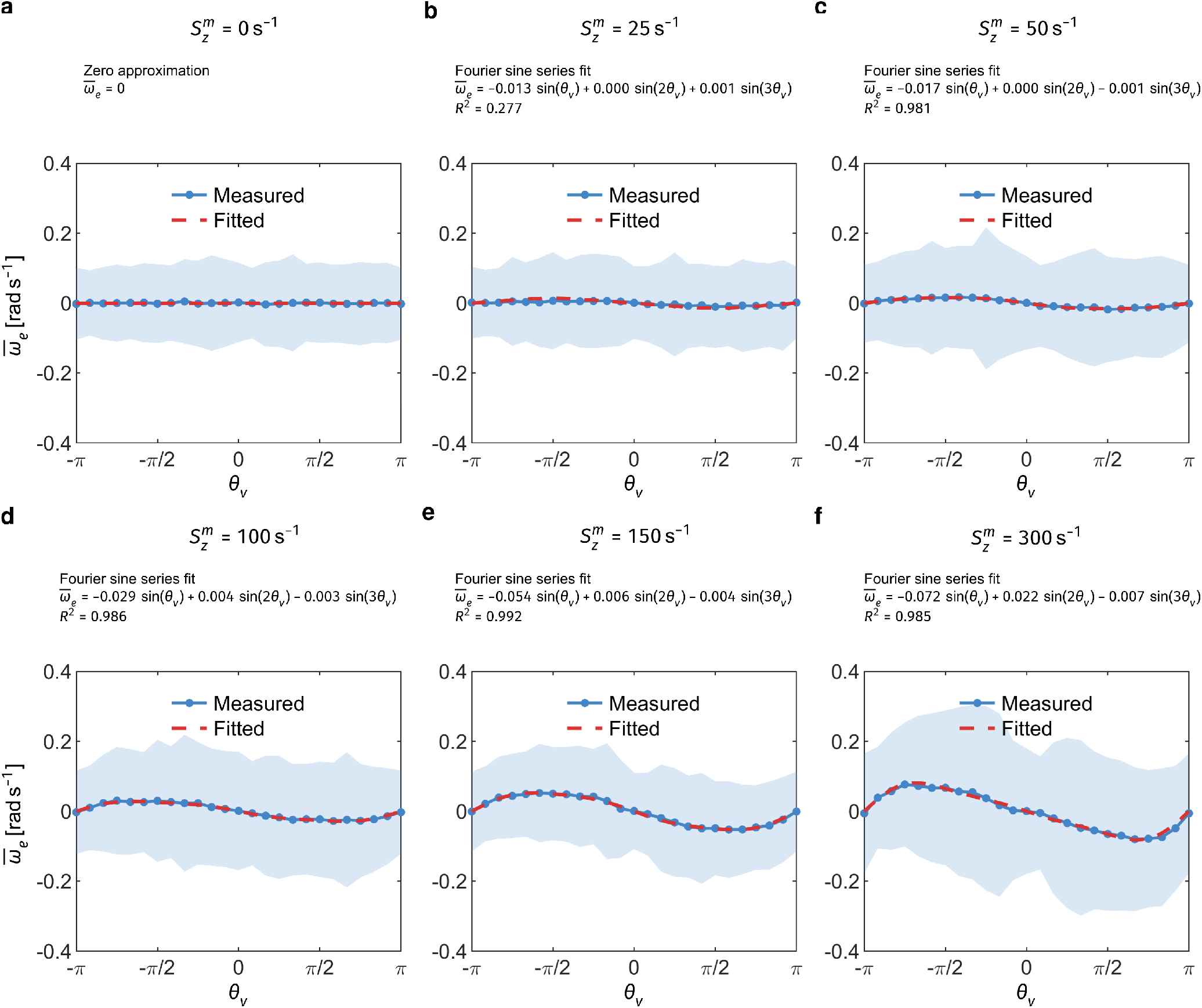
Mean angular velocity 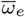 as a function of direction *θ*_*v*_ at different bed shear rates. For 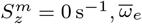 is approximately zero. For 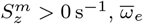 is represented by a third-order Fourier sine series. **a**–**f**, 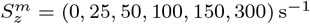.

**Extended Data Fig. 7.**
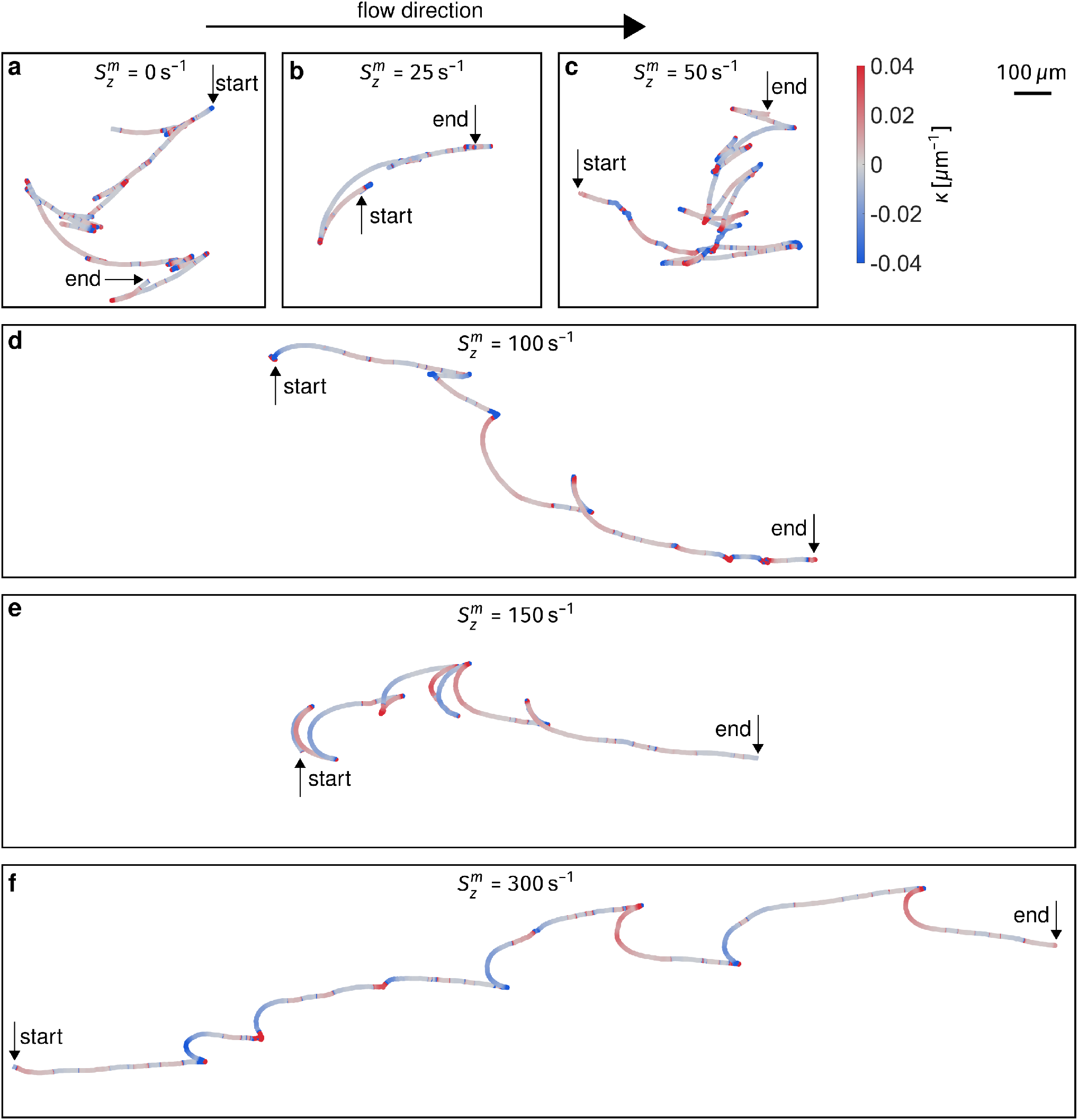
Representative trajectories at all tested bed shear rates, coloured by path curvature. **a**–**f**, 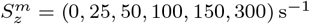.

**Extended Data Fig. 8.**
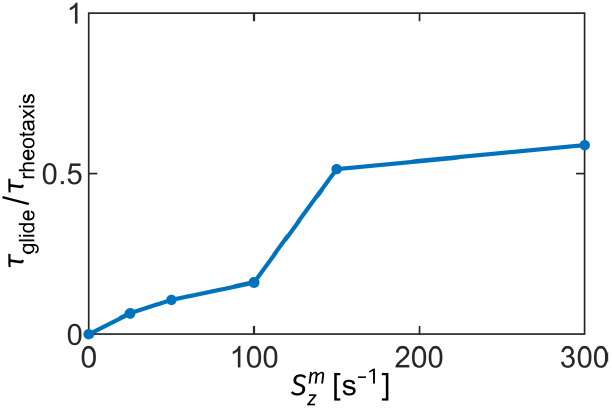
Ratio of the timescale for a continuous gliding event to that for shear-induced reorientation at different bed shear rates. The continuous gliding timescale, *τ*_glide_, is taken as the mean dwell time of the gliding motility state shown in Extended Data Fig. 3. The shear-induced reorientation timescale, *τ*_rheotaxis_, is defined as the time required for reorientation from *θ*_*v*_ = 3*π/*4 to *θ*_*v*_ = *π/*4, based on the fitted angular velocity at different bed shear rates shown in Extended Data Fig. 6.

**Extended Data Fig. 9.**
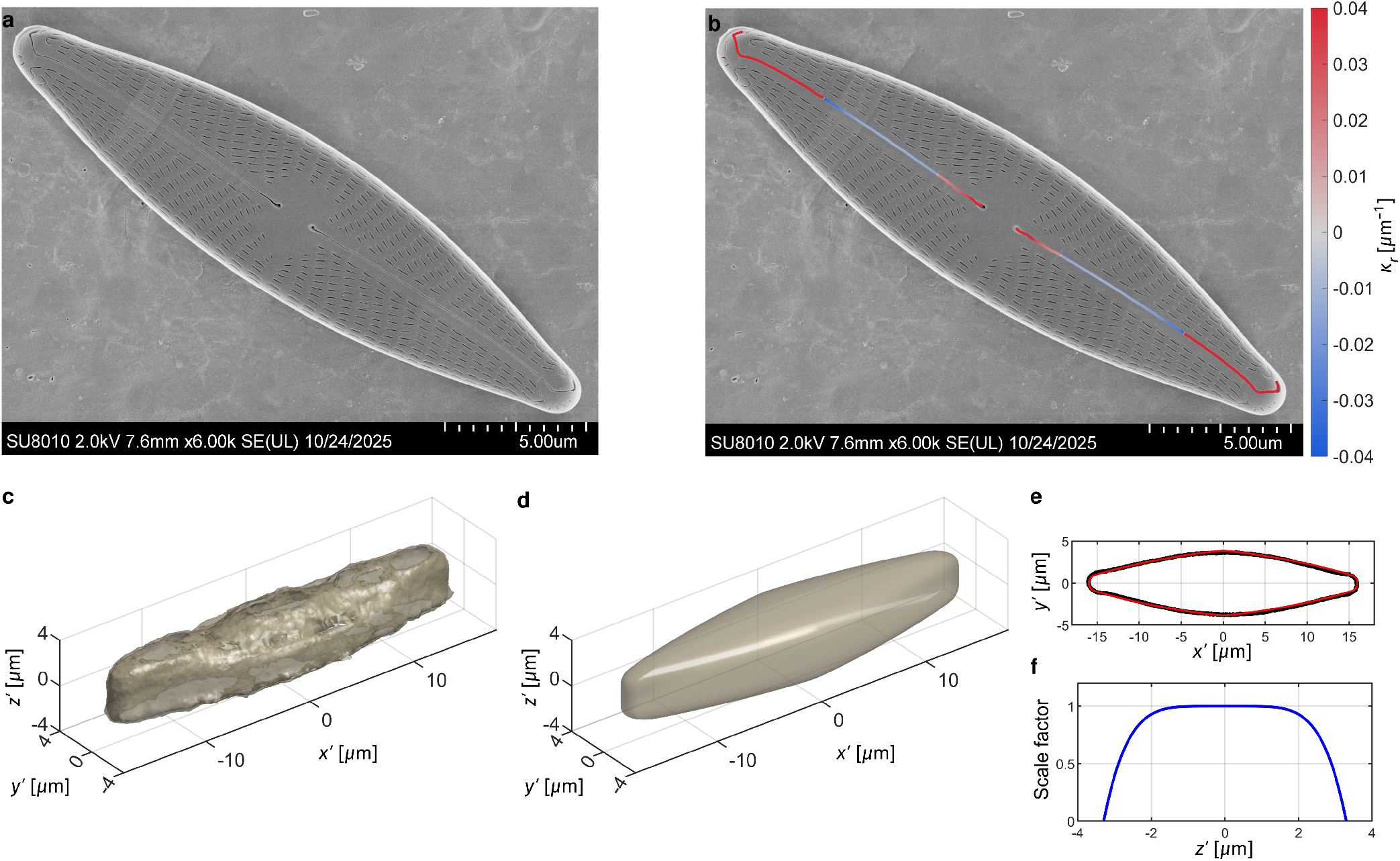
SEM images and 3D reconstruction of *N. cryptocephala* from z-stack imaging. **a**, Original image of the upper valve face. **b**, Estimated raphe curvature *κ*_*r*_ along the two raphe branches separated by the central nodule. **c**, Raw z-stack scan of *N. cryptocephala*. **d**, Reconstructed 3D model based on the z-stack scan. **e**, Fit of the maximal plane from the z-stack scan. The red line indicates the fitted curve, and the black dots indicate the raw data. **f**, z-dependent length scale factor of the maximal plane for reconstructing the model from the z-stack scan. In **c**–**f**, *x*^′^, *y*^′^, and *z*^′^ denote the coordinates along the major axis, minor axis, and vertical axis, respectively, with the origin at the cell centre.

**Extended Data Fig. 10.**
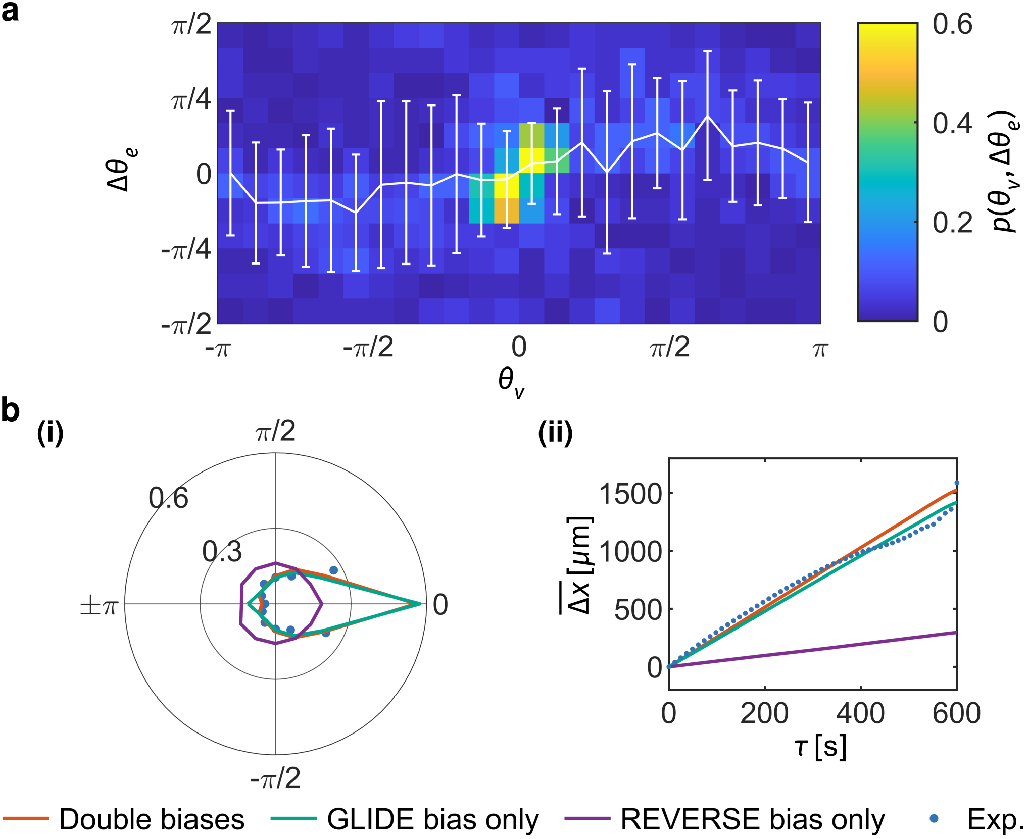
Sensitivity of simulated rheotaxis to angular biases in the GLIDE and REVERSE states. **a**, Joint probability density of the gliding direction *θ*_*v*_ before the REVERSE event and the change in long-axis orientation across the REVERSE event, Δ*θ*_*e*_. White line and error bars indicate the mean and standard deviation of Δ*θ*_*e*_ in each *θ*_*v*_ bin. Under shear, REVERSE events exhibit a weak downstream bias: after polarity switching, cells tend to emerge with a small angular offset whose sign reduces the absolute angular deviation of the gliding direction after the REVERSE event from the downstream direction. This tendency is consistent with both individual trajectories (Extended Data Fig. 7) and the population-level statistics shown here. **b**, Comparison of stochastic-model predictions at 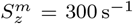 for different combinations of angular bias. (**i**), Distribution of gliding direction *θ*_*v*_ . (**ii**), Mean streamwise drift 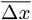 as a function of lag time. ‘Double biases’ denotes the full model including both the downstream-restoring angular velocity during GLIDE and the downstream-biased angular offset during REVERSE. ‘GLIDE bias only’ retains only the GLIDE-state angular bias, whereas ‘REVERSE bias only’ retains only the REVERSE-state angular bias. The GLIDE bias alone already reproduces the measured rheotactic alignment and downstream drift quantitatively and remains close to both the model with double biases and experiment. By contrast, the REVERSE bias alone yields substantially weaker rheotaxis and deviates strongly from both the full model and the data. Thus, the REVERSE-associated bias makes only a secondary contribution, mainly by further suppressing the upstream-oriented peak after REVERSE, whereas the dominant origin of downstream rheotaxis is the shear-induced angular dynamics during GLIDE.

