## Supplementary Information for "Benthic diatoms navigate shear flows via hydrodynamic rolling and active gliding"

### Supplementary Videos

**Supplementary Video 1.** Bright-field imaging of *Navicula cryptocephala* motility at $S_z^m = 300\,\mathrm{s}^{-1}$. The movie is shown at 60× real speed.

**Supplementary Video 2.** Cell trajectories extracted from bright-field imaging in still water. The current cell positions are indicated by white ellipsoids, and previous trajectories are shown as magenta solid lines. No obvious directional bias is observed. The movie is shown at 60× real speed.

**Supplementary Video 3.** Cell trajectories extracted from bright-field imaging at $S_z^m = 100\,\mathrm{s}^{-1}$. The current cell positions are indicated by white ellipsoids, and previous trajectories are shown as magenta solid lines. Pronounced downstream migration is observed. The movie is shown at 60× real speed.

**Supplementary Video 4.** Cell trajectories extracted from bright-field imaging at $S_z^m = 150\,\mathrm{s}^{-1}$. The current cell positions are indicated by white ellipsoids, and previous trajectories are shown as magenta solid lines. Pronounced downstream migration is observed. The movie is shown at 60× real speed.

**Supplementary Video 5.** Cell trajectories extracted from bright-field imaging at $S_z^m = 300\,\mathrm{s}^{-1}$. The current cell positions are indicated by white ellipsoids, and previous trajectories are shown as magenta solid lines. Pronounced downstream migration is observed. The movie is shown at 60× real speed.

**Supplementary Video 6.** Interference reflection microscopy imaging of cell–substrate contact dynamics at $S_z^m = 300\,\mathrm{s}^{-1}$. The cell maintains a stable downstream gliding direction. The frustule outline is shown as a magenta dashed line, the contact site is marked by a red cross, and the previous trajectory is shown as a magenta solid line. The movie is shown at real speed.

**Supplementary Video 7.** Interference reflection microscopy imaging of cell–substrate contact dynamics at $S_z^m = 300\,\mathrm{s}^{-1}$. The cell initially glides upstream in an unstable direction. The frustule outline is shown as a magenta dashed line, the contact site is marked by a red cross, and the previous trajectory is shown as a magenta solid line. The movie is shown at real speed.

**Supplementary Video 8.** Interference reflection microscopy imaging of cell–substrate contact dynamics at $S_z^m = 300\,\mathrm{s}^{-1}$. The cell starts with a unstable gliding direction perpendicular to the flow and soon reorients downstream. The frustule outline is shown as a magenta dashed line, the contact site is marked by a red cross, and the previous trajectory is shown as a magenta solid line. The movie is shown at real speed.

**Supplementary Video 9.** Interference reflection microscopy imaging of cell–substrate contact dynamics in still water. The frustule outline is shown as a magenta dashed line, the contact site is marked by a red cross, and the previous trajectory is shown as a magenta solid line. The estimated contact site exhibits random shifts along the longitudinal axis on either side of the cell centre, while the lateral displacement remains small. The movie is shown at real speed.
